# Mitochondrial efficiency determines Crabtree effect across yeasts

**DOI:** 10.1101/2024.11.01.621473

**Authors:** Julius Battjes, Pranas Grigaitis, Milou Hoving, Thomas D. Visser, Karina Stampfl, Gabriela A. Miguel, Johan van Heerden, Jesper K. Andersen, Frank J. Bruggeman, Bas Teusink

## Abstract

Under excess glucose conditions, many yeasts switch from high-yield respiratory metabolism to low-yield fermentation, a phenomenon called the Crabtree effect in yeast, or the Warburg effect in mammalian cells. Cellular constraints and limited resources are generally believed to govern the metabolic strategies of cells to adapt to environmental conditions, but which constraints drive this switch is still under debate. Here we study the Crabtree-negative, fully respiratory yeast *Pichia kluyveri* and compare it to the Crabtree-positive yeast *Saccharomyces cerevisiae* from a resource allocation perspective. By integrating quantitative physiology and proteomics into whole-cell proteome-constrained models, we find that the Crabtree effect is determined by the composition and catalytic efficiency of the electron transport chain. We find that the subsequent proteome efficiency of respiration versus fermentation varies between these species. The variation in parameters and composition of the respiratory machinery likely reflects the evolutionary and ecological history of these yeast species. This study advances our understanding of the role of proteome constraints and proteome efficiency in governing cellular metabolism of yeasts, and that of eukaryotic cells at large.

## Introduction

Overflow metabolism, also called the (bacterial) Crabtree effect or Warburg effect, is the phenomenon that under conditions of nutrient excess or fast growth, cells switch from full respiratory metabolism with a high yield of ATP on sugar, to a lower-yield metabolic mode, such as fermentation. This occurs under fully aerobic conditions and is a widely occurring metabolic phenomenon in both bacterial, yeast, and mammalian cells [1–3].

The Crabtree effect is often studied experimentally in a chemostat [2]. Following the example of yeasts, Crabtree positive *S. cerevisiae* switches metabolism as a response to changing dilution rate (which in steady state equals the growth rate) in a chemostat. *S. cerevisiae* fully respires in cultures below so-called critical growth rate, and increasingly switches to ethanol fermentation upon increasing dilution rate, either linearly [4] or in two phases [5, 6]. This phenomenon has been subject of many studies, as it appears counter-intuitive that a high ATP-yielding strategy (respiration) is traded for a low yield strategy (fermentation) at high growth rates where ATP demand is high [7].

Numerous hypotheses have been proposed to explain the onset of overflow metabolism in aerobic environment. The predicted behavior following these hypotheses, switching to the overflow phenotype, typically coincides with the outcome of a constrained optimisation problem with growth rate as a proxy for fitness [8, 9]. Different formulations of the optimization problem have in common that use of low substrate yield pathways is predicted as the outcome by additional active constraint(s) that forces cells to increase growth rate at the expense of this substrate yield (reviewed in [10]). Therefore, not substrate yield but ATP yield on protein (proteome efficiency) explains the optimal growth maximizing strategy in nutrient-rich conditions (see, e.g., [6, 11, 12]). Knowledge of this type of metabolic reprogramming in various growth strategies is of great importance in both biomedical and biotechnological research and applications.

The availability of advanced metabolic models and multi-omics data has enabled formulation of new theories on underlying reasons for the switch. Recently, we have investigated the constraints that govern the Crabtree effect in *S. cerevisiae* with a so-called proteome-constrained model, or pc-model [6, 13]. With these models, we can investigate in great detail how resources are optimally allocated under specific conditions.

We found that the active constraints during the metabolic shift from respiration to respiro-fermentation depended on the conditions: at the onset of ethanol formation in glucose-limited chemostats, the (experimentally determined) upper bound on the mitochondrial space became an active constraint, but at even higher growth rates - at glucose excess - the constraint on total proteome capacity was active. The latter means that all available protein space was occupied by growth-supporting proteins and therefore, any reallocation of proteome space to further increase in growth rate would come at the expense of other proteins. Therefore, the fact that respiration was actively repressed in favor of fermentation under these conditions, and thus that an increase in growth rate was possible by a trade-off between respiration for fermentation, must mean that fermentation is more “proteome efficient” - in this yeast.

The concept of proteome efficiency, however, has different definitions in the literature, and this has caused some confusion. In a recent paper, Shen et al. used derived ATP flux over invested protein as the definition of proteome efficiency, as also used by Basan et al. for *Escherichia coli* [12, 14]. Shen et al. concluded that respiration is always more proteome efficient than respiration, and attributed the Crabtree effect to adaptation of *S. cerevisiae* to anaerobic growth, rather than optimal resource allocation. Deriving efficiency metric from observed proteome allocation means that the efficiency is now condition-dependent and describes the actual state, and not the potentially optimal state, and hence does not explain the Crabtree effect from some first (evolutionary) principles or prevailing active constraints. Basan et al. used the catalytic rate normalized by molecular weight as the protein cost, which is condition-independent but has a disadvantage that the cost is not directly expressed in terms of the growth rate increase. Moreover, both methods described above require to define the set of proteins that belong to respiration or fermentation, and these sets are not trivial to define [15]. Our protein-constrained models compute the pathway that leads to the highest growth rate under proteome constraints, and thus computes by definition the most proteome efficient route without having to define catabolism and anabolism *a priori*.

The yeast phylum is diverse in Crabtree phenotypes [16]; an useful comparison to explain the Crabtree effect in yeast is to compare a Crabtree positive yeast to a Crabtree-negative one under similar conditions and with the same type of analysis. Here we studied Crabtree-negative yeast *Pichia kluyveri* (*P. kluyveri*), which is used for its preferred respiratory metabolism in the production of nonalcoholic and low-alcohol beverages, but is relatively unexplored when it comes to quantitative physiology [17]. We first established the Crabtree-negative phenotype in aerobic glucose excess conditions. Next, we built an extensive dataset by quantifying metabolic fluxes, biomass composition, proteome composition, and available volumetric proteome space of *P. kluyveri* in both carbon-limited and carbon-excess conditions. Finally, we modified the existing pcYeast framework to represent the metabolism of *P. kluyveri* (pcPichia) and integrated the experimental data to parametrize the pcPichia model. By thorough comparison between a Crabtree-positive and Crabtree-negative yeast, we identified the factors that drive this metabolic difference.

## Results

### 0.1 *P. kluyveri* is Crabtree negative

We started with characterizing *P. kluyveri* by growing on different carbon, nitrogen, and vitamin sources. This was followed by a thorough physiological characterization in carbon-limited and nutrient excess (batch) conditions (Supplementary Data 1). We quantified multiple physiological metrics: exometabolite fluxes, biomass composition, protein expression, cell size, and mitochondrial area (Fig. 1 A). Furthermore, we sequenced the (industrial) strain and reconstructed the first genome-scale metabolic model (GEM) of *P. kluyveri* from its sequenced genome and curated the GEM with the obtained characterization (see Supplementary Notes for details).

**Fig. 1.**
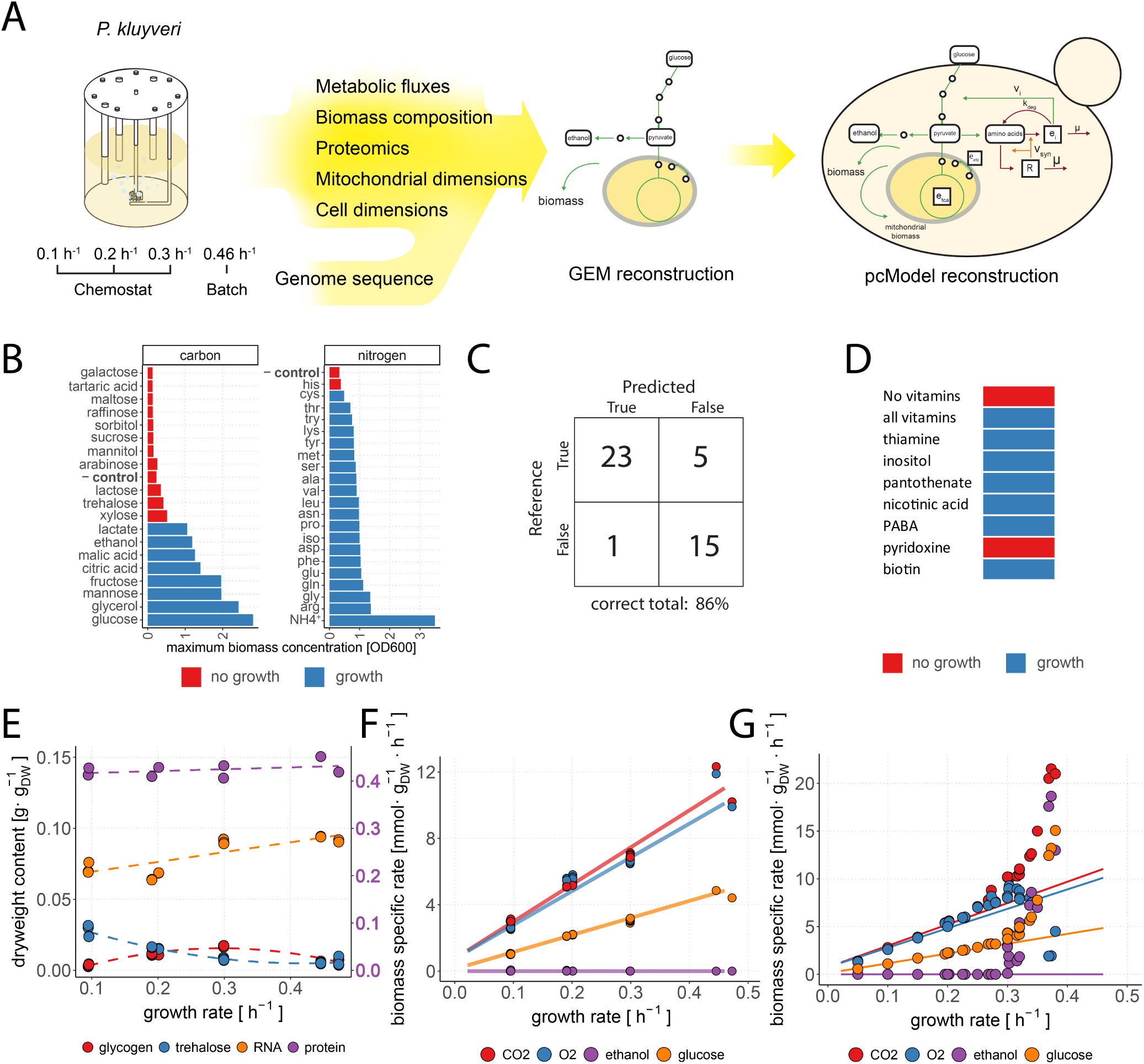
Physiological characterization of *P. kluyveri* and GEM reconstruction. (A) Schematic overview workflow used in this study. (B) Growth evaluation on different C- and N-sources. We considered *P. kluyveri* to grow on a carbon or nitrogen source if there was significantly more (p>0.05, independent t-test) biomass formation compared to negative control (media without carbon or nitrogen source). (C) Confusion matrix of growth predictions by GEM for growth on C- and N-sources. (D) Results of vitamin drop-out experiment which led to identification of pyridoxine auxotrophy for *P. kluyveri*. (E) Biomass content measurements for glycogen, trehalose, RNA, (left y-axis) and protein (right y-axis) from cultures in glucose-limited and excess conditions. (F) Biomass-specific consumption and excretion rates (fluxes) of *P. kluyveri* in glucose-limited and excess conditions. We used our curated GEM of *P. kluyveri* to predict metabolite fluxes, shown in solid lines. (G) GEM prediction data for *P. kluyveri* in lines overlaid with *S. cerevisiae* data in similar conditions [4, 6].

*P. kluyveri* was unable to grow on oligosaccharides like maltose and trehalose as carbon sources, as previously reported [18] (Fig. 1 B). We observed growth on urea, ammonia, and all amino acids except histidine as the sole nitrogen source. We used the GEM to predict which C- and N-sources *P. kluyveri* can use, and predictions were correct in 86 % of the cases (Fig. 1 C). In addition, we found that pyridoxine biosynthesis pathway in the *P. kluyveri* GEM was incomplete, with pyridoxal-5’-phosphate synthase (SNZ1) missing in genome. *P. kluyveri* thus is auxotrophic for pyridoxine (Fig. 1 D).

Next, we grew *P. kluyveri* in bioreactors under glucose-limited and glucose-excess conditions to determine metabolic fluxes and biomass composition of *P. kluyveri* (Table 1). Measurements included protein, RNA, glycogen, and trehalose biomass content. Protein content remained relatively stable at 41 % across growth rates; RNA content of *P. kluyveri* increased with increasing growth rate (Fig. 1 E). In contrast, the trehalose content decreased with the growth rate and the glycogen content was highest at 0.3 *h*^−1^. We updated the biomass equation of *P. kluyveri* GEM using this data to obtain a more realistic representation (Supplementary Figure 1; Supplementary Data 2; see Supplementary Notes for details).

**Table 1.** Physiological parameters and specific rates of *P. kluyveri* in glucose excess, glucose limited, and ethanol excess conditions. Biomass specific rates (*q*) are given in 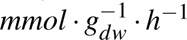. Biomass yield on glucose (*Y_x/s_*) is given in [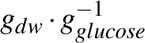].

| carbon source | glucose | glucose | glucose | glucose | ethanol |
| --- | --- | --- | --- | --- | --- |
| growth rate [$h^{-1}$] | 0.1±0 | 0.2±0.01 | 0.3±0 | 0.46±0.02 | 0.31±0.02 |
| $q_{CO2}$ | 3.01±0.06 | 5.18±0.09 | 6.9±0.13 | 11.27±1.5 | 8.43±1.09 |
| $q_{O2}$ | 2.58±0.06 | 5.6±0.11 | 6.68±0.16 | 10.9±1.4 | 16.73±2.22 |
| $q_{acetate}$ | 0.02±0.01 | 0.03±0 | 0.04±0.01 | 0.13±0.01 | 0±0 |
| $q_{ethanol}$ | 0.02±0.02 | 0.01±0.02 | 0±0 | 0±0 | 9.74±0.27 |
| $q_{glucose}$ | 1.03±0.03 | 2.15±0.05 | 3.06±0.08 | 4.64±0.31 | 0±0 |
| $q_{glycerol}$ | 0±0 | 0±0 | 0±0 | 0.06±0.02 | 0±0 |
| $q_{pyruvate}$ | 0±0 | 0±0 | 0±0 | 0.01±0 | 0±0 |
| $Y_{x/s}$ | -0.51±0.01 | -0.5±0 | -0.54±0.02 | -0.55±0.06 | -0.69±0.03 |
| carbon balance [%] | 108.27±3.04 | 97.94±0.77 | 99.61±3.14 | 104.66±4.17 | 103.71±6.79 |

No metabolic shift was evident from measured metabolic fluxes in glucose-limited and excess experiments; with increasing growth rate up to *µ_max_*, *q_o_*_2_, *q_co_*_2_, and *q_glucose_* showed a linear increase with growth rate (Fig. 1 E). In addition, no major carbon fluxes were missing as carbon balances closed around 100 %. In all conditions we observed fully respiratory growth, with the exception of a minor acetate production flux at *µ_max_* (with flux 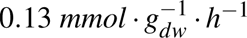)(Table 1). The biomass yield (*Y_x/s_*) of *P. kluyveri* varies between 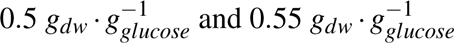 (Table 1). Overall, we conclude that *P. kluyveri* is completely respiratory and therefore Crabtree-negative in the measured conditions.

With measurements of both the major metabolic fluxes and biomass composition, we parametrize the GEM, i.e. to determine energy maintenance coefficients of *P. kluyveri*. Therefore, both non-growth-(NGAM) and growth-associated maintenance (GAM) were fitted to match experimental fluxes of *P. kluyveri* in glucose-limited and excess conditions. Predicted match measured fluxes with an NGAM of 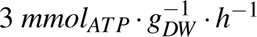 and GAM of 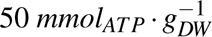 (Fig. 1 E).

The biomass-specific consumption and production rates of *P. kluyveri* are comparable to respiratory *S. cerevisiae* (before the onset of fermentation) (Fig. 1 G). After the onset of fermentation, around 0.27 *h*^−1^, *S. cerevisiae* switches to fermentative metabolism, with a maximum growth rate of 0.41 *h*^−1^. In contrast, *P. kluyveri* remains completely respiratory and does not show signs of switch metabolic strategy up until the maximum growth rate of 0.46 *h*^−1^. This superior respiratory metabolism is also reflected in its growth on ethanol as carbon source, with a growth rate of 0.31 *h*^−1^ (Table 1) as compared to 0.15 *h*^−1^ in *S cerevisiae* [19]. In contrast to *S cerevisiae*, *P. kluyveri* cannot grow anaerobically (results not shown). We therefore confirmed that *P. kluyveri* is a fully aerobic, Crabtree-negative yeast with a distinct flux profile from *S. cerevisiae*.

### 0.2 Protein expression is subject to both metabolic and hierarchical control

Next we checked how the proteome of *P. kluyveri* aligns with this physiological characteristic. We performed label-free mass spectrometry-based proteomics analyses on glucose-limited and excess condition samples (Fig. 2). In individual runs, we quantified between 3219 and 4639 proteins, or from 46 % to 65 % of 7180 predicted proteins in the annotated proteome of *P. kluyveri*. To enable functional interpretation of the proteomics data, we manually curated the protein, pathway, and sub-cellular localization annotation of > 80 % of the expressed proteins (Supplementary Data 3).

**Fig. 2.**
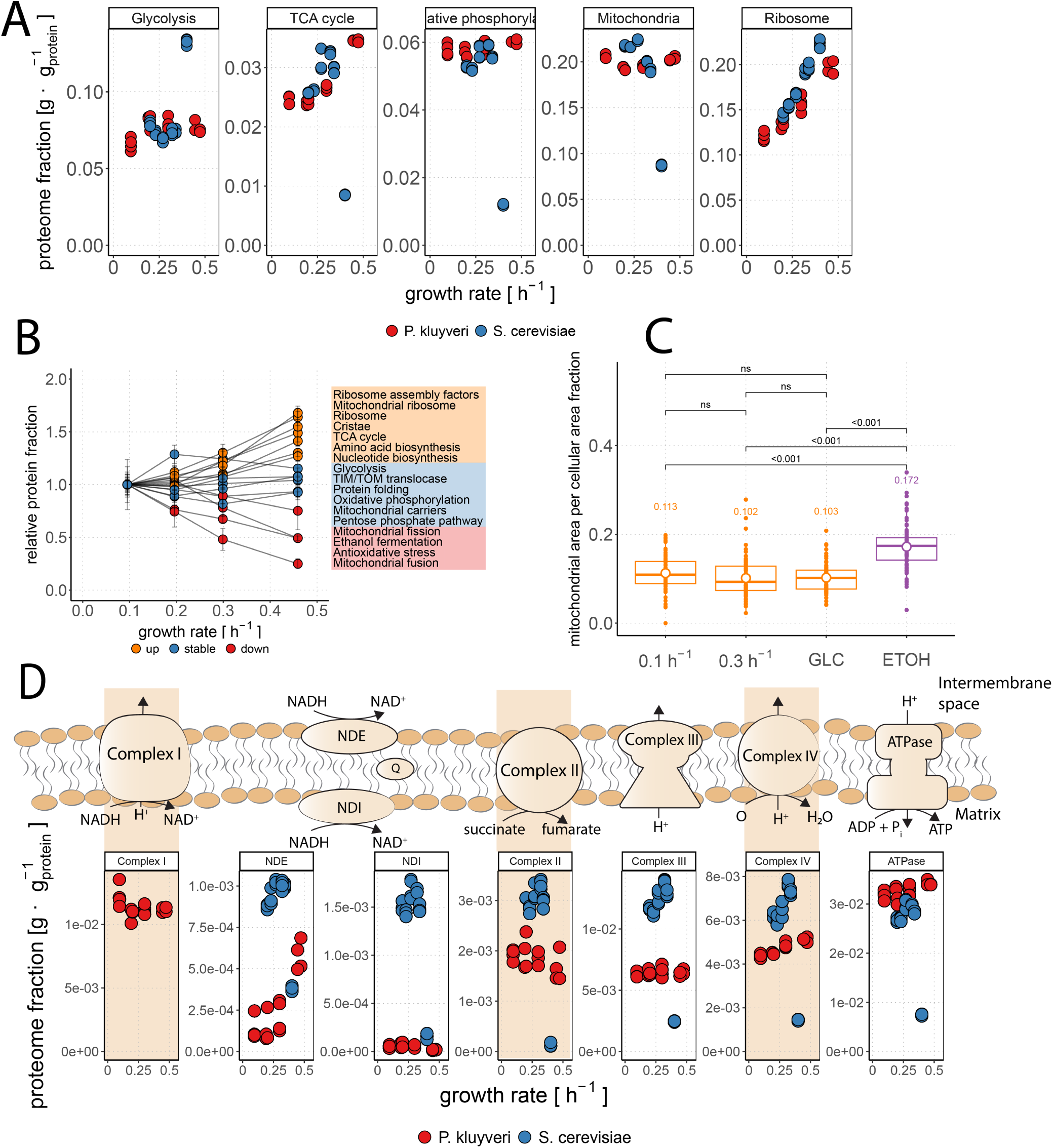
Proteome analysis of *P.kluyveri* compared to *S. cerevisiae* (A) Protein expression levels of both yeasts lumped into pathways (Supplementary Data 3). (B) Protein expression levels of pathways to relative to the reference condition of *µ* = 0.1 for *P. kluyveri*. (C) Fraction of relative mitochondrial area per cellular area in single *P. kluyveri* cells. Area was determined from segmented EM pictures. Relative area was determined for aerobic glucose limited (0.1*h*^−1^ & 0.3*h*^−1^), glucose batch (GLC), and ethanol batch (ETOH) conditions. (D) Expression levels of individual oxidative phosphorylation proteins in *P. kluyveri* and *S. cerevisiae* with schematic representation of oxidative phosphorylation. All proteome data of *S. cerevisiae* is taken from Elsemman et al. [6].

While all major metabolic fluxes (*q_o_*_2_, *q_co_*_2_, and *q_glucose_*) showed a linear increase with growth rate (Fig. 1 A), we observed only minor changes in pathway expression levels of TCA cycle, oxidative phosphorylation, glycolysis and mitochondria in glucose-limited cultures, and even at the maximal growth rate under glucose excess (Fig. 2 A). This observation is contrary to previously acquired *S. cerevisiae* proteomics data [6]. As *µ* → *µ_max_*, the proteome of *S. cerevisiae* shifts to a more fermentative expression profile, with major rearrangement in the proteome: fold-increase in glycolytic protein levels and decreasing protein levels in the TCA cycle, oxidative phosphorylation, and consequently decreasing total mitochondrial proteome.

In contrast to catabolic pathways, we observed a more uniform increase in expression of anabolic pathways, including ribosomes, amino acid biosynthesis, and nucleotide biosynthesis (Fig. 2 A & B). In addition, the proteome fraction of ribosomal proteins linearly increased with increasing growth rate, just as in *S. cerevisiae* (Supplementary Figure 4). In contrast to *S cerevisiae*, ethanol fermentation-related genes showed decreasing expression with increasing growth rate, even though no ethanol production was measured in any of the conditions (Fig. 2 B).

Limited space of mitochondrial proteome was identified as a determinant of the onset of ethanol fermentation at the critical growth rate (*µ_crit_*) in glucose-limited chemostats for *S. cerevisiae* [6]. Current published data on mitochondrial capacity are limited to measurements in *S. cerevisiae* [20–23] (Supplementary Table 1). Therefore, we measured the mitochondria-to-cell ratio at different growth rates (and carbon sources) for *P. kluyveri* by the relative mitochondrial surface area for *P. kluyveri* using electron microscopy (EM) (Supplementary Data 4). Mitochondrial area does not differ significantly and remained around 10 − 11 % for glucose grown cultures (Fig. 2 C). This is in agreement with our constant mitochondrial proteome data under the same conditions (Fig. 2 A). In ethanol batch cultivations, the mitochondrial content increased to 17 %, hinting at a higher mitochondrial requirement of growth on ethanol. This is more than double the observed mitochondrial content of 7.6 % in ethanol batch cultivations of *S. cerevisiae* [21].

In summary, we observed hierarchical control on anabolic pathways of *P. kluyveri*, as protein expression increased with growth rate. In contrast, we saw largely metabolic control on catabolic pathways, where expression levels of oxidative phosphorylation remained relatively constant, independent of growth rate and hence flux.

### 0.3 Mitochondria of *P. kluyveri* harbor functional respiratory Complex I

We then zoomed in at the expression of proteins involved in the electron transport chain and oxidative phosphorylation. We identified most subunits for Complex I in the genome of *P. kluyveri*, which were also found expressed in the proteomics data set (Fig. 2 D). In *P. kluyveri* multiple routes of electron transfer are possible: An alternative to Complex I is internal NADH dehydrogenase (NDI) that passes electrons from mitochondrial matrix NADH onto ubiquinone, or external NADH dehydrogenase (NDE), which transfers electrons from cytosolic NADH.

The use of Complex I within the ETC has implications for the ATP yield on substrate in *P. kluyveri*. For example, *S. cerevisiae* does not have a lot of flexibility in its electron transport chain; it does not have Complex I but uses NDI and reaches a P/O (*ATP* · *O*^−1^) ratio of 0.95 [24]. With this P/O ratio, theoretical ATP yield on glucose 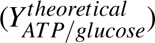 is around 16 *ATP* · *glucose*^−1^ in *S. cerevisiae*. Unlike NDI, Complex I contributes to the proton motive force (pmf) by pumping protons from the mitochondrial matrix into mitochondrial intermembrane space, with a stoichiometry of 4 protons per reduced NADH [25]. We used the constructed GEM of *P. kluyveri* to determine the subsequent P/O ratio of oxidative phosphorylation, which increased from 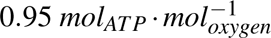 with NDI to 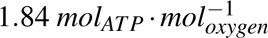 with Complex I. In turn, Complex I increases 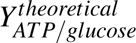 from 15.4 *ATP* · *glucose*^−1^ to 23.4 *ATP* · *glucose*^−1^ (see Supplementary Notes for detailed discussion).

We found that the expression of each protein complex remained relatively stable, including Complex I (Fig. 2 D). The gene annotated as NDE (2172_g) increased expression levels with increasing growth rate, which fits with its role to donate biomass-formation-associated cytosolic NADH to the ETC. NDI (3903_g) on the other hand decreased its (very low) expression levels; both NDE and NDI levels were substantially lower than Complex I (Fig. 2 D). From the proteome data we conclude that *P. kluyveri* primarily used Complex I to reoxidize NADH in these conditions.

### 0.4 Proteome efficiency explains Crabtree effect

Next, we wanted to explore the impact of the observed differences in physiology and proteome allocation between Crabtree negative and positive yeast. Therefore, we developed a pc-model for *P. kluyveri*, as these models compute to the best of our abilities the integrated resource costs of metabolic growth strategies, and the active cellular constraints (see Supplementary Notes for details) (Supplementary Figure 2, Supplementary Data 5).

In pc-models, we predict the minimal protein level by setting the upper bound of fluxes to the *V_max_* = *k_cat_* · [*e*] of the enzyme. Subsequently, by comparing the minimal-vs. measured protein levels, we can infer the apparent saturation of enzymes or pathways. If minimal level predictions were higher than measured protein levels, however, this indicates that for these enzymes the *k_cat_*values must be too low. Therefore, we systematically increased the *k_cat_* values of such enzymes such that at maximal flux (near *µ_max_*) the predicted minimal enzyme level was equal to the measured proteome fraction. This was needed for several reactions in the electron transport chain and TCA cycle. We defined this adjusted value as the effective *k_cat_* (*k_cat,e_*) (Table 3). In contrast, we found higher expression levels than minimally required at full saturation for glycolytic enzymes, as we observed and discussed before for *S. cerevisiae* [6, 26] (Fig. 3; Supplementary Figure 7). We will further refer to the pc-models with updated *k_cat,e_*values as effective.

**Fig. 3.**
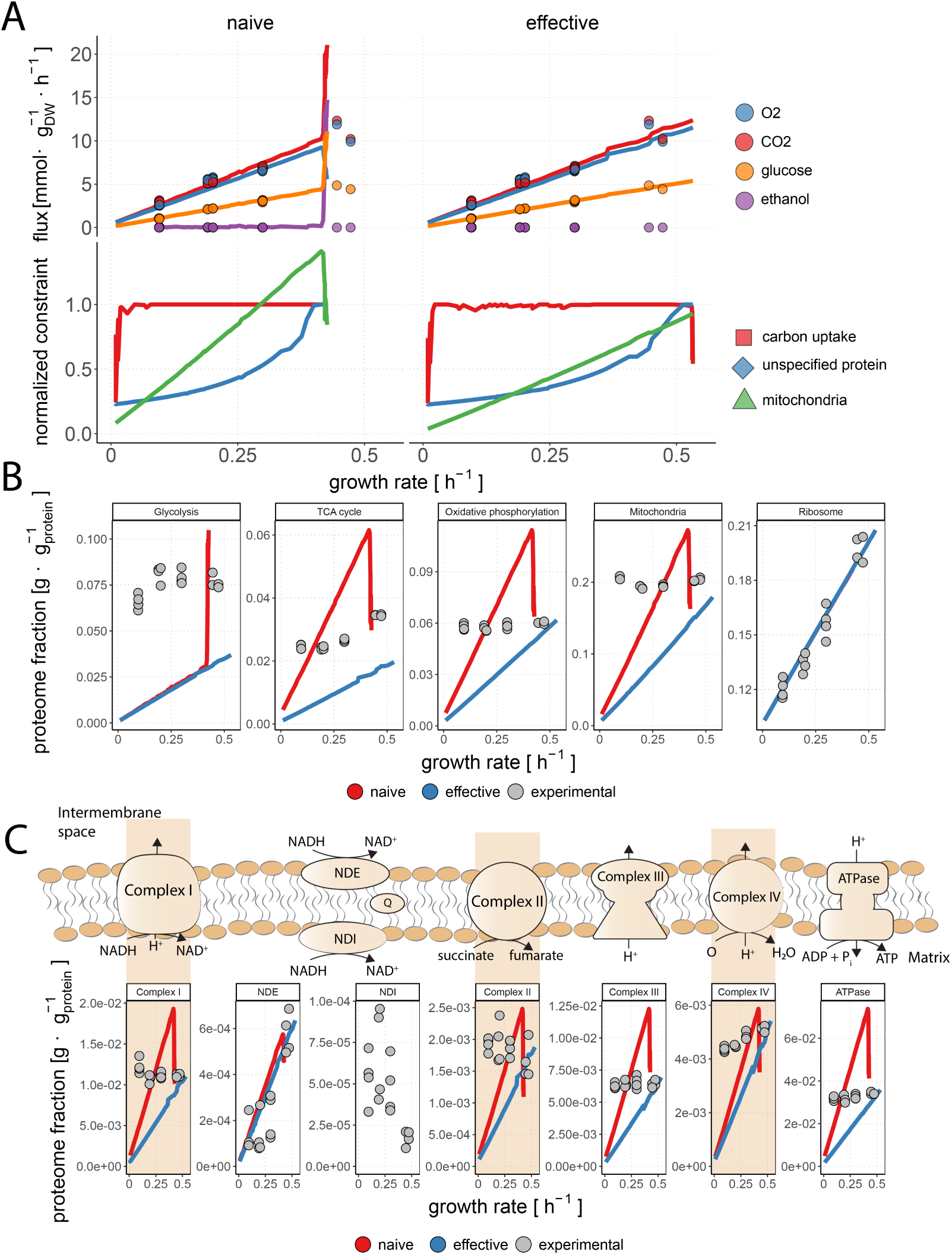
Proteome efficiency determines pcPichia preference for respiration (A) (top) Flux profiles of pcPichia (lines) and experimental *P. kluyveri* data of glucose limited chemostat and glucose batch experiments (points). In pcPichia (naïve), no catalytic constants were modified with respect to pcYeast (naïve). In pcPichia (effective), catalytic constants were changed by comparison between pc-model predictions and measured proteome (Table 3). (bottom) Constraint evaluation of pcPichia for total proteome, plasma membrane, and mitochondrial volume constraints. The expression of the constraint equals one when the maximal capacity of the compartment is reached, and therefore the compartment capacity limits growth. (B) Predicted expression levels of proteins in central metabolic pathways and mitochondria compared to measured proteome fractions. (C) Expression levels of individual oxidative phosphorylation proteins for the different pcYeast configurations.

To compare *P. kluyveri* with *S. cerevisiae*, we first redid simulations of the pcYeast model (Supplementary Figure 7 A). We determined *k_cat,e_*’s for TCA cycle and oxidative phosphorylation for pcYeast (effective) by comparing predicted protein levels to measured protein levels at *µ_crit_*, where respiratory protein expression levels are highest (Supplementary Figure 7 B & C, Supplementary Figure 6 B). In pcYeast we limited how much mitochondrial (protein) volume fits in the cell with a mitochondrial volume constraint that was estimated from orthogonal data [6]. Once the mitochondrial volume constraint is hit, oxidative phosphorylation is limited and ethanol fermentation starts to cover increased ATP demands. After adjusting catalytic constants in pcYeast (effective), we needed to adjust this mitochondrial volume constraint to set the critical growth rate to the experimentally observed 0.28*h*^−1^ in pcYeast (effective). Therefore, as in pcYeast, as we increase glucose availability still first mitochondrial volume constraint is hit and fermentation starts. Subsequently, when *µ* → *µ_max_*the total proteome constraint was hit and fermentation flux was further increased and respiratory protein was suppressed. Therefore, after adjusting catalytic constants, fermentation remained more proteome efficient in *S. cerevisiae*.

We used our experimental measurements to parametrize pcPichia (Table 2, see Supplementary Notes). We first simulated growth in glucose-limited chemostats at increasing dilution rate using the naïve pcPichia model, which is the model with kinetic parameters based on *S. cerevisiae* and the GECKO method [27]. We identified two different growth regimes (Fig. 3 A): At low D, the constraint on the expression level of glucose transport was the only active constraint and the model was respiratory (Fig. 3 B). Now, we also set an upper bound to how many protein fit in the cell, derived from protein biomass content measurements. We observed that with increasing glucose availability and hence growth rate, this total proteome constraint was hit near *µ_max_*and respiratory protein was exchanged for fermentative protein (Fig. 3 B & C). The maximal growth rate that the model could reach was lower than experimentally observed. The fact that the model shifts from respiration to fermentation to increase growth rate when the total proteome constraint becomes active suggests that fermentation is more proteome efficient than respiration, by definition, even in this model where Complex I is present.

**Table 2.** Comparison of parameters used in the pc-model formulations of *P. kluyveri* (pcPichia) and *S. cerevisiae* (pcYeast) [13]. How minimal non-growth associated protein (NGAP) fraction of the cell (*NGAP_min_*), mitochondrial NGAP (*NGAP_min,mitochondrial_*) (Supplementary Figure 2 C), ribosomal fraction at zero growth 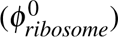, ribosomal elongation rate (*k_elongation_*) (Supplementary Figure 4), cell volume (*V_cell_*)(Supplementary Figure 5), and mitochondrial available cell volume (*V_mitochondrialprotein_*) are determined is discussed in Supplementary Notes.

| parameter | *P. kluyveri* | *S. cerevisiae* |
| --- | --- | --- |
| Complex I | yes | no |
| PO ratio [$ATP \cdot O^{-1}$] | 1.825 | 0.95 |
| $NGAP_{min}$ [$g \cdot g_{protein}^{-1}$] | 0.19 | 0.22 |
| $NGAP_{min,mitochondrial}$ [$g \cdot g_{mitochondrialprotein}^{-1}$] | 0.14 | 0.11 |
| $GAM$ [$mmol_{ATP} \cdot g_{dw}^{-1}$] | 40 | 24 |
| $NGAM$ [$mmol_{ATP} \cdot g_{dw}^{-1} \cdot h^{-1}$] | 0.5 | 0.168 |
| $\phi_{protein}$ [$g_{protein} \cdot g_{dw}^{-1}$] | 0.42335 | $0.42436 \cdot \mu + 0.35364$ |
| $V_{cell}$ [$\mu m^3$] | 22.7 | $29.3716 \cdot \mu + 20.55066$ |
| $\phi_{ribosome}^0$ [$g \cdot g_{protein}^{-1}$] | 0.1 | 0.053 |
| $k_{elongation}$ [$aa \cdot s^{-1}$] | 16 | 10.5 |
| $\Psi_{mitochondria}$ [$\mu m_{mitochondrial}^3 \cdot \mu m_{cell}^3$] | 0.106 | 0.074 |
| $V_{mitochondrialprotein}$ [$\mu m^3$] | 0.791 | 0.7 |

We then updated *k_cat_*’s to match minimal predicted protein levels to experimentally observed ones. We also adjusted the mitochondrial volume constraint of pcPichia based on the mitochondrial volume constraint of *S. cerevisiae*, using an experimentally derived conversion factor between relative mitochondrial volumes of *P. kluyveri* and *S. cerevisiae* (see Supplementary Notes for details). PcPichia (effective) predicted no metabolic shift and no ethanol production in glucose-limited- and glucose-excess conditions, consistent with the Crabtree-negative physiology of *P. kluyveri* (Fig. 3 A). Moreover, the maximal growth rate on glucose was now more in line with experimental data. PcPichia (effective) also correctly predicted fluxes in ethanol batch when corrected for non-growth associated protein content (Supplementary Figure 3). The total proteome constraint still was the only active constraint at *µ_max_*, but fermentation was no longer favored over respiration. In pcPichia (effective), we observe that at *µ_max_*, the mitochondrial volume constraint and total proteome constraint converge, yet the mitochondrial volume constraint is never reached (Fig. 3 A).

Therefore, when Complex I is present and the efficiency of respiration is increased according to proteomics data, respiratory metabolism becomes more proteome-efficient than fermentation. Below *µ_max_*, model predictions underestimate the required amount of protein, indicative of the flux through the electron transport chain (and the TCA cycle) being regulated at the metabolic level, not at the level of protein expression.

### 0.5 Composition of electron transport chain and catalytic constants determine efficiency of catabolism

Our experimental and model analysis suggests that *P. kluyveri* maintains a Crabtree-negative phenotype due to interplay of two factors: higher ATP yield from respiration (P/O ratio), and higher catalytic constants. Harboring only one of them, e.g., presence of Complex I alone is not sufficient (Fig. 3). When comparing different yeast species, we see that there is a heterogeneity when it comes to presence of the Crabtree effect and ETC composition (Table 4). Consequently, we investigated which parameters in our model could potentially influence the switching between Crabtree-positivity and -negativity. We started by removing Complex I from pcPichia with updated *k_cat_* values (pcPichia (effective, no Complex I)) and observed that respiration remains favourable (Fig. 4 A). In contrast, when adding Complex I to pcYeast (effective + Complex I), respiration becomes more proteome efficient (Fig. 4 B). Fermentation still starts when mitochondrial constraint is hit but respiration is no longer exchanged when total proteome constraint is hit.

**Fig. 4.**
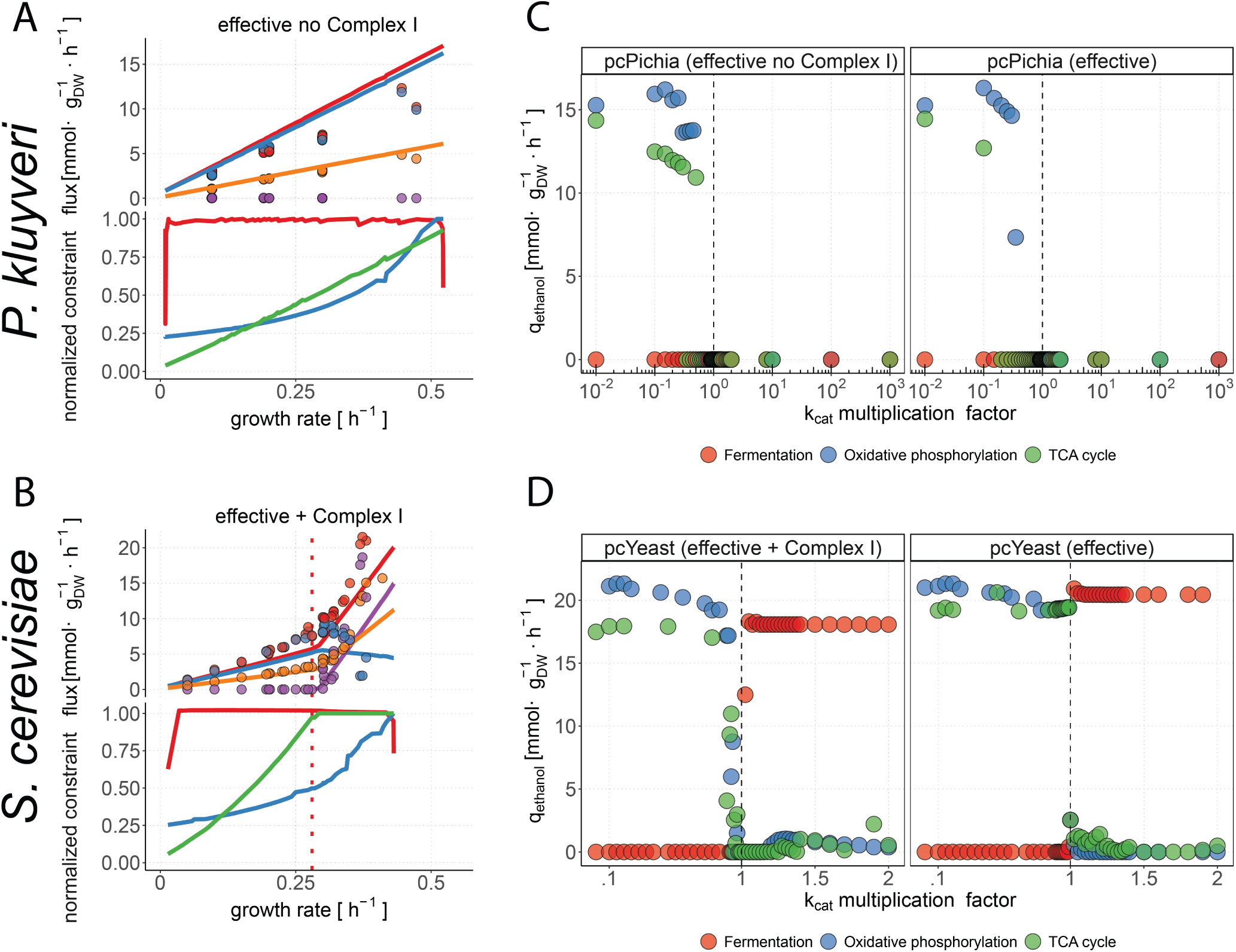
Change of individual catalytic constants is enough to change the more proteome efficient metabolic strategy. (A) (top) Flux profiles of pcPichia (effective, without Complex I) (lines) and experimental *P. kluyveri* data of glucose limited chemostat and glucose batch experiments (points). (bottom) Constraint evaluation of pcPichia for total proteome, plasma membrane, and mitochondrial volume constraints. The expression of the constraint equals one when the maximal capacity of the compartment is reached, and therefore the compartment capacity limits growth. (B) Same as in A but for pcYeast (effective, with Complex I). (C) The ethanol flux at *µ* → *µ_max_*as a function of catalytic constants of proteins in TCA cycle, oxidative phosphorylation, and fermentation. We titrated the the catalytic rate constants *k_cat_* and observed switching from fully respiratory to fermentative metabolism. Here, predictions are shown for pcPichia (effective) both with (left) and without (right) Complex I. (D) same as in (C) but for pcYeast (effective) with (left) and without (right) Complex I.

**Table 3.**
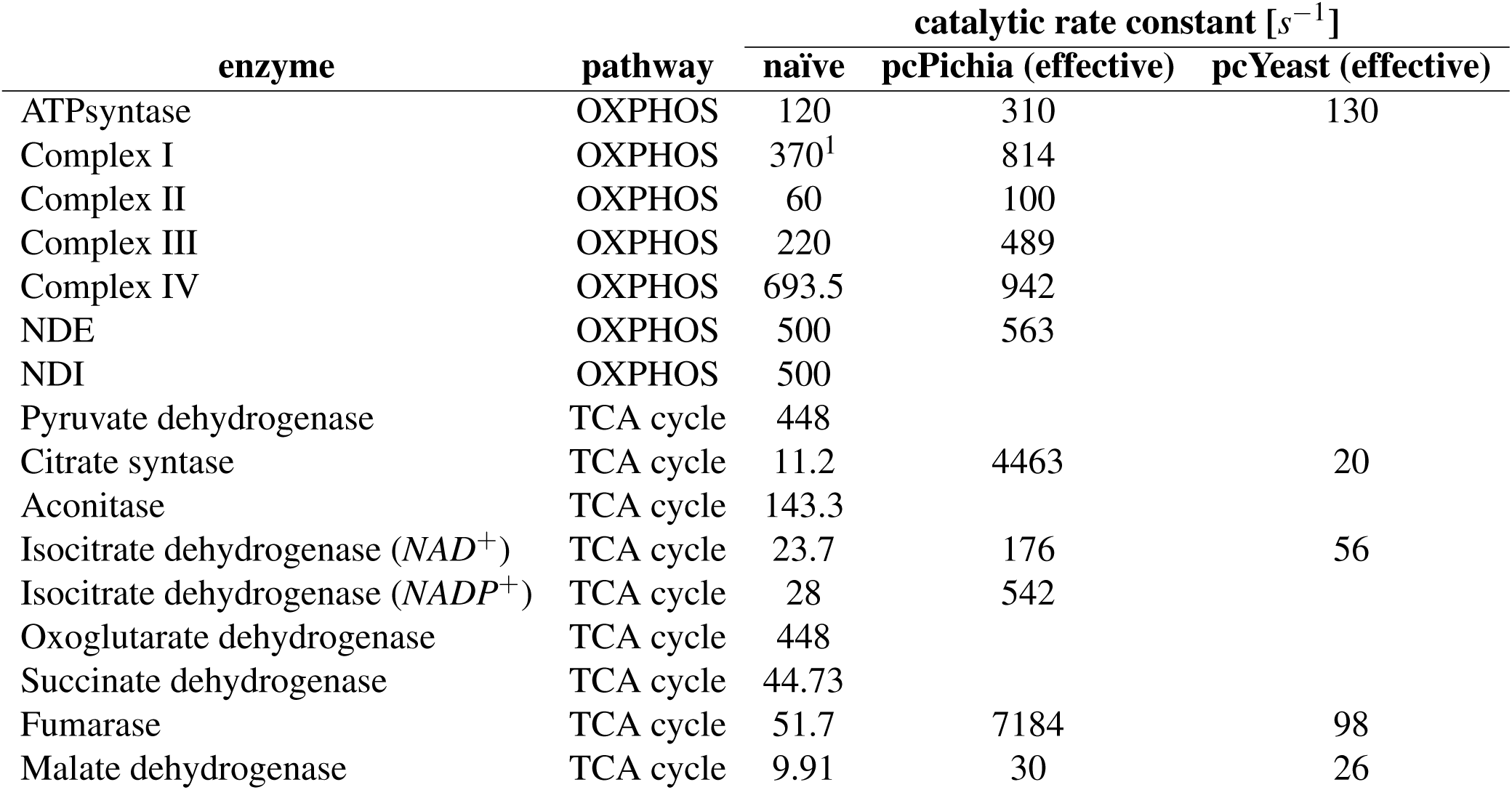
Manually curated catalytic rate constants for respiratory chain- and TCA cycle proteins. *k_cat_* for naïve models are given as used in Grigaitis et al. (2023) [13]. 1. [57]. A more complete overview of model parameters for each model can be found in Supplementary Data 5.

**Table 4.** Diversity of the composition of electron transport chain and Crabtree phenotype across different yeasts.

| **species** | **Complex I** | **NDE / NDI** | **Crabtree** | **reference** |
| --- | --- | --- | --- | --- |
| *S. pombe* | F | T | Positive | [58, 59] |
| *K. marxianus* | F | T | Negative | [58] |
| *L. thermotolerans* | F | T | Positive | [58] |
| ***P. kluyveri*** | T | T | Negative | this study |
| *S. cerevisiae* | F | T | Positive | [2] |
| *D. bruxellensis* | T | T | Positive | [58] |

Now that we have observed that catalytic constants can vary between the two yeast species (Table 3), we tested whether the catalytic constants for the oxidative phosphorylation, TCA cycle and fermentation change the predicted phenotype, i.e. changes the proteome efficiency differences in favor of either respiration or fermentation. We also tested how the presence of Complex I influenced the outcome of pcPichia (effective) and pcYeast (effective). When removing Complex I in pcPichia(effective), we needed to decrease the catalytic constants of all protein in oxidative phosphorylation or TCA cycle by a factor around 0.5, for pcPichia to become fermentative at *µ_max_* (Fig. 4 C). Substantially lower catalytic constants were required to become fermentative in the presence of Complex I. Increasing only the catalytic constants of fermentation (pyruvate decarboxylase and alcohol dehydrogenase) did not result in fermentation.

For pcYeast we removed mitochondrial constraint for this analysis, and therefore ethanol production is now only present when fermentation has a higher proteome efficiency. We observe a switch for both pcYeast models, with or without Complex I, with relative small changes in catalytic constants for both TCA cycle, oxidative phosphorylation, and fermentation (Fig. 4 D). With a slight increase in catalytic constant for oxidative phosphorylation or TCA cycle pcYeast without Complex I becomes respiratory and with a slight decrease pcYeast with Complex I becomes fermentative. In a similar fashion, decreasing catalytic constant of fermentation without Complex I results in higher proteome efficiency for respiration, while increasing catalytic constant with Complex I results in fermentation.

We conclude that resource allocation theory can explain the Crabtree negative physiology of *P. kluyveri* as the interplay of electron transport chain composition and catalytic constants. The difference in the Crabtree effect between *S. cerevisiae* and *P. kluyveri* can be attributed to the difference in proteome efficiency between the two yeast species, which is largely caused by the expression of Complex I, but also the exact *k_cat_* values play an important role.

## Discussion

Many hypotheses exist about the molecular- and economical principles of why respiration-fermentation switches happen in some organisms, yet the question of what the underlying determinants are, remains open. The Crabtree effect is not a universal trait among yeasts, and therefore we could look for explanations inside this microbial phylum; here we extensively characterized the Crabtree negative metabolism of *P. kluyveri* and compared its physiology to the Crabtree positive yeast *S. cerevisiae*.

We grew *P. kluyveri* in glucose-limited conditions at different dilution rates and confirmed that *P. kluyveri* is Crabtree-negative (Fig. 1 E & F). We found that the fluxes of the two yeast species were strikingly similar in the respiratory regime of *S. cerevisiae*, before the onset of fermentation at critical growth rate. This is surprising in view of the higher ATP yield in oxidative phosphorylation for *P. kluyveri* due to Complex I, whose proteins were consistently expressed under all conditions examined (Fig. 2 D).

We found that use of Complex I is preferred over NDI in our resource allocation models despite substantial investments into biosynthesis, transport, and maintenance of Complex I subunits (a total of 1067 kDa per functional complex) versus the NDI bypass (a single protein of 76 kDa). The difference between investments associated with these two NADH reoxidation routes inspired discussions whether exploiting NDI, and not Complex I, could be a valid strategy in growth on glucose, as experimentally shown for the yeast *Ogataea parapolymorpha* [28, 29]. However, our proteomics data supports use of Complex I over NDI - although we can also not exclude that Complex I is expressed but not used. However, to arrive at a yield on substrate *Y_x/s_* comparable to that of respiratory *S. cerevisiae*, a higher maintenance energy expenditure was needed to consume the surplus ATP of *P. kluyveri*’s higher-yield oxidative phosphorylation. An explanation for this observation remains an open question beyond the scope of this paper, and the origin(s) of maintenance energy expenditure remains a topic of debate [15, 28].

In addition to electron transport chain composition, we found that catalytic constants play a crucial role in determining the Crabtree phenotype. For *P. kluyveri*, we changed catalytic constants in both TCA cycle and oxidative phosphorylation to match protein level predictions at *µ_max_* with experimental data, which resulted in a higher proteome efficiency of respiration (Fig. 3 B & C). We showed that when these catalytic constants are changed collectively, the more efficient metabolic strategy can shift from fermentation to respiration and vice versa. For *P. kluyveri* Complex I presence increased the required factor to change the more proteome efficient route and therefore plays a major but not a necessarily decisive role in determining the Crabtree effect. For *S. cerevisiae* changes in relative proteome efficiency were already observable in a narrow range of 5% of the used catalytic constants.

For *S. cerevisiae* the critical dilution rate for onset of ethanol formation is not dependent on proteome efficiency [6, 30, 31]. It seems that for *S. cerevisiae* mitochondrial volume is a regulatory constraint; here we imposed this constraint to obtain a critical growth rate at 0.28*h*^−1^ for the updated pcYeast model. Intriguingly, *P. kluyveri* does not hit this mitochondrial volume constraint until *µ_max_*, where it converges with total proteome constraint. We corrected the *P. kluyveri* mitochondrial volume constraint for difference in relative mitochondrial volumes between *P. kluyveri* and *S. cerevisiae* (see Supplementary Notes for details). Notably, we found no data for mitochondrial volume fraction in *S. cerevisiae* around *µ_crit_*. Although there is a 30 % difference in mitochondrial volume fraction between the two species, we observe little difference between mitochondrial proteome fractions while respiratory (Fig. 2 A). When the mitochondrial volume constraint is left uncorrected, *P. kluyveri* hits the mitochondrial volume constraint before reaching *µ_max_*, which results in ethanol fermentation as *µ* → *µ_max_* (Supplementary Figure 8). Since we do not observe this experimentally, it appears that mitochondrial volume fractions must differ between the species even while the mitochondrial proteome fractions are similar, suggesting a difference in mitochondrial proteome density between *P. kluyveri* and *S. cerevisiae*. Alternatively, metabolic regulation of the mitochondrial enzymes differ and the *k_cat_*’s of respiratory enzymes in *P/ kluyveri* are higher. This calls for a more detailed comparative study on the physiology and morphology of the mitochondria in these two species.

*S. cerevisiae* is respiratory when glucose transport limited growth as respiration has the highest ATP on substrate yield, but respiration is exchanged for fermentation when growth is proteome limited - showing that fermentation is more proteome efficient. *P. kluyveri* is respiratory with Complex I when glucose transport limited and remains respiratory when proteome limited as this pathway has highest ATP on substrate yield and is most proteome efficient. Among the parameters that distinguish the two yeast species other than the mitochondrial ones, we found that *P. kluyveri* has a higher ribosome elongation rate, a characteristic that often varies among different yeast species (Table 2) [32]. Considering resource allocation, a higher ribosome elongation rate will not benefit proteome efficiency of fermentation more than respiration, but contributes the higher maximal growth rate observed in *P. kluyveri*.

If overflow metabolism is indeed linked to optimal resource allocation towards growth of microbes, it is subject to the precise accounting of the interplay between returns (yield of ATP per NADH reoxidized) and investments (proteome efficiency) of the catabolic machinery, the respiratory chain in particular. This may explain the difficulty of finding one overarching explanation for the absence or presence of the Crabtree effect in yeasts: the economy of the cell is shaped by the ecological and evolutionary history of the yeasts, and then we come into tricky waters. The lack of solid kinetic, proteomic, and physiological data across many organisms brings us to speculation; for instance, Shen et al. [14] proposed that adaptation to anaerobic conditions may explain the preference for fermentation for *S. cerevisiae*. Yet, if this was true, another Crabtree-positive yeast *Schizosaccharomyces pombe* has collected all the wrong cards: *S. pombe* exhibits a higher P/O ratio of 1.28 compared to *S. cerevisiae*, but does not grow well anaerobically [33, 34]. It appears therefore that we have not yet reached the rock bottom of the evolutionary origin of the Crabtree effect.

To conclude, we present a detailed description of the industrially relevant Crabtree negative yeast *P. kluyveri*. We show that using knowledge on *S. cerevisiae* in combination with precisely targeted measured data on *P. kluyveri* we can model its Crabtree negative physiology. Finally, we show that when growth is proteome constrained, metabolic strategies are governed by the efficiency of proteins, which in turn are influenced by ecological factors.

## Methods

### Media, strains, and maintenance

*P. kluyveri* NEER was obtained from Novonesis A/S (Hørsholm, Denmark). *S. cerevisiae* CEN.PK-113.7D was obtained from P. Kötter, Euroscarf (Frankfurt, Germany). Cells were stored in −80 ^◦^*C* in 30 %(*w/v*) glycerol. Yeast from glycerol stocks was grown on solid YP2%D (10 *g* · *l*^−1^ Bacto yeast extract, 20 *g* · *l*^−1^ Bacto peptone, 20 *g* · *l*^−1^ agar, and 20 *g* · *l*^−1^ D-glucose) agar plates before further cultivation. Shake flask growth experiments were performed at 30 ^◦^*C* whilst shaken at 200 rpm in an orbital shaker in synthetic media (SM), unless stated otherwise [24].

### DNA extraction

Cells were cultivated in 5 *g* · *L*^−1^ yeast extract (Oxoid, England), 10 *g* · *L*^−1^ neutralized soya peptone (Oxoid, England), 10 *g* · *L*^−1^ glucose (Sigma-Aldrich, Germany) at pH 5.6. Culture was grown overnight at 25 ^◦^*C* with orbital shaking at 100 rpm. The cell pellet was harvested by centrifuging at 4000 x g, 4 ^◦^*C* for 15 minutes and washed with 30 *mL* of PBS buffer (*pH* 7.0).

High molecular DNA from *P. kluyveri* was isolated using Qiagen Genomic Tip 100/g (Qiagen, Germany) following manufacturer’s instructions. To preserve high molecular size of the DNA, use of vortex was avoided along the entire protocol. Quality of the isolated DNA was assessed as follows: DNA was quantified with Invitrogen Qubit™ Fluorometer, double-stranded DNA (dsDNA) Broad Range (ThermoFisher, USA) according to manufacturer’s instructions; DNA molecular size was assessed with 5200 Fragment Analyzer System with the kit HS Genomic DNA 50 kb (DNF-468-0500) (Agilent Technologies, USA) and the software ProSize 3.0 (Agilent Technologies, USA); DNA purity was assessed through the spectrophotometer Nanodrop™ (Thermo Fisher, USA).

### Library preparation

DNA libraries for Nanopore sequencing (Oxford Nanopore Technologies, UK) were prepared according to the manufacturer’s protocol using the Ligation Sequencing Kit SQK-LSK109 (Oxford Nanopore Technologies, UK). In brief, 2 *µg* of DNA was diluted with 49 *µL* nuclease-free water and DNA was end-repaired and dA-tailed using NEBNext® Ultra™ II End Repair/dA-Tailing Module according to the manufacturer’s protocol (New England Biolabs Inc., USA). Cleaning was done by adding 1:1 of beads to the samples and the beads were washed with 200 *µL* of ethanol 70 %. DNA was eluted in 25 *µL* of nuclease-free water and quantified with Invitrogen Qubit™ Fluorometer as described above. For the native barcode- and the adapter ligation, the kit EXP-NBD104 and EXP-NBD114 600 (Oxford Nanopore Technologies, UK) and the reagents Blunt/TA Ligase Master Mix, NEBNext Quick Ligation Reaction Buffer (5X) and Quick T4 DNA Ligase (New England BioLabs Inc., USA) were used following instructions from the manufacturers. The pooled libraries were diluted to 65 *µL* of nuclease-free water. The samples were incubated at 38 ^◦^*C* for 30 minutes to increase the recovery of long DNA fragments. Afterwards, magnetic beads were pelleted and the clear eluate was retained in a new tube. The finished libraries were quantified as described with Invitrogen Qubit™ Fluorometer as described above.

Genomic libraries for Illumina sequencing were generated using NEBNext® Ultra™ II FS DNA Library Prep Kit for Illumina® with NEBNext Multiplex Oligos for Illumina (Unique Dual Index UMI Adaptors DNA Set 1), (New England Biolabs Inc., USA) on Biomek i5 Liquid Handler (Beckman Coulter, USA). 200 *ng* of genomic DNA diluted in 15 *µL* EB buffer (Tris-Cl, *pH* 8.0) was used in the half-volume reaction mixes for fragmentation, end-repair/A-tailing, ligation and final amplification. Fragmentation time was optimized to 8 minutes. 5 *µL* of 2.5 *µM* NEBNext Multiplex Oligos for Illumina (Unique Dual Index UMI Adaptors DNA Set 1), (New England Biolabs Inc., USA) was used during adapter ligation step. 10 *µL* of the adapter-modified DNA fragments were enriched by 9-cycle PCR. Clean NGS beads (Clean NA, The Netherlands) were used for double-sided post ligation size selection and one post-amplification clean-up to purify fragments at average size between 450 to 550 bp. Concentration of gDNA and double stranded DNA libraries were measured by QubitFlex® Fluorimeter using Qubit dsDNA Broad range and Qubit 1x dsDNA HS assays (Thermo Fisher Scientific, USA), respectively. Average dsDNA library size distribution was determined using the Agilent HS NGS Fragment (1-6000 bp) kit on the Agilent Fragment Analyzer (Agilent Technologies, USA). Libraries were normalized and pooled in the normalization buffer (10 *mM* Tris-Cl, *pH* 8.0, 0.05 % Tween 20) to the final concentration of 10 *nM*.

### Sequencing

Long read sequencing was performed in a Flow Cell R9.4.1 in MinIon device (Oxford Nanopore Technologies, UK) with MinKNOW software v2.0 (Oxford Nanopore Technologies, UK) following the manufacturer’s protocol. The estimate N50 was 20 *Kb*. Basecalling was performed using Guppy v5.0.11. For short read sequencing, 1 *pM* pool of libraries denaturated in 0.2 *M* NaOH and 1300 *µL* ice-cold HT1 buffer was loaded onto the flow cell provided in the NextSeq Reagent Mid Output (300 cycles) kit and sequenced on a NextSeq 550 platform (Illumina, USA) with a paired-end protocol and read lengths of 151 *nt*.

### Hybrid de novo assembly

Long reads were trimmed for size and identity using Filtlong v0.2.0 (Wick, 2017) with the parameters –min_length 1000 –keep_percent 90 and –target_bases 500000000. Hybrid *de novo* genome of *P. kluyveri* was assembled using Canu v2.0 [35] with the parameters –genomeSize=12m and –nanopore-raw. Draft assembly was polished with the short reads using the software Pilon v.1.22 [36]. Functional annotation of the assembled *P. kluyveri* NEER genome was performed with eggNOG-mapper v2 using default annotation options [37].

### Bioreactor cultivation

Bioreactor cultivations were performed in 1.5L reactors (Applikon, Delft) with 1 L working volume. Cultivations were stirred at 600 rpm and temperature was maintained at 30 ^◦^*C*. The pH was maintained at 5.0 (AppliSens, Applikon) with 2 M KOH. SM media was used with 0.5 *g* · *l*^−1^ glucose end concentration for both glucose excess and limited conditions. Dissolved oxygen levels remained above 40 % and were analyzed by Clark electrodes (AppliSens, Applikon). The bioreactor was controlled with ez2Control controllers (Applikon, Delft) in combination with software from Lucullus (Securecell BV, Rotterdam). From inoculation to the first sample (batch) or till turning on the pumps (chemostat) ambient air was used for cultivation. For the remainder of the cultivation time, compressed air was used at 0.6 *l* · *min*^−1^. In chemostat cultivations, a steady state was assumed after nine generation times, of which fermentation parameters (metabolite concentrations, biomass dry weight, and both oxygen and carbon dioxide concentrations) remained stable.

### Gas analysis

A condenser cooled the off-gas from the reactor. Oxygen and carbon dioxide content was measured online with a Servomex ME4100 (Servomex, Zoetermeer). Calibration of the Servomex gas analyzer was performed with pure nitrogen gas and a mixture of 20.8 % O2 and 3 % CO2 (Nippon Gases, Dordrecht).

### Bioreactor sampling

For metabolite analysis of the broth, broth was filtered with 0.22 *µm* PES filter (Whatman). For steady-state chemostat samples, the broth was first quenched with cold steel beads followed by PES filter filtration [38]. For determination of internal metabolite concentrations of glycogen and trehalose cells were processed according to the leakage-free rapid quench method [39]. Biomass was harvested by centrifugation for 10 minutes at 4000 rpm and 4 ^◦^*C*, and washed twice in either ice-cold phosphate buffer saline (PBS)(pH 7.4) or DEPC-treated demi water for RNA quantification samples. Biomass pellets were snap-frozen in liquid nitrogen and stored at −80 ^◦^*C*. Mitochondrial staining was performed on the second pre-steady state sample of chemostat experiments. Culture was diluted in respective media type to final OD600 between 0.5 and 1.0 Transcription was stopped by the addition of 0.125 *mg* · *mL*^−1^ cycloheximide [40].

### Analytic methods

Organic acid concentrations were determined by high performance liquid chromatography (HPLC) analysis on a Prominence HPL chromatograph (SHIMADZU, Kyoto, Japan) equipped with a Rezex ROA organic acid H+ column (Phenomex, California, USA), with *H*_2_*SO*_4_ as a running fluid. Glucose concentration of chemostat samples was determined by enzymatic assay. In short, in a flat bottom 96 wells plate, 25 *µl* of sample was mixed with reaction mixture (PIPES buffer (30.22 *g* · *l*^−1^, pH 7.0), NADP (4.5 *mM*), ATP (4.5 *mM*) and *MgSO*_4_ (5 *mM*)). Blank absorbance was subsequently measured at 340 *nm*. Thereafter 10 *µl* of hexokinase/ glucose-6-phosphate dehydrogenase (Roche, Mannheim, Germany) was added, and the endpoint was measured again at 340 *nm*.

OD600 of cultures was measured with an Ultraspec 2100 pro (Amersham Biosciences, Cambridge, England). Biomass dry weight was measured on 2 *µm* pre-weighed membrane filters (Whatman). Filters were subsequently washed with demi water after which the filter was dried for at least 24 hours at 60 deg*C* before weighing.

### Biomass content measurements

Glycogen and trehalose content of biomass were determined by enzymatic degradation to glucose of by amyloglucosidase (Sigma) and trehalase (Sigma), respectively. This degradation to glucose was followed by enzymatic quantification of glucose [41]. Protein content in biomass was determined by copper sulphate reaction in 1M NaOH with bovine serum albumin (BSA) as a standard [24]. RNA content of biomass was measured by cold-acid extraction followed by the modified Benthin quantification protocol [42, 43]. 0.02 %(*v/v*) Tyloxapol was added to the 0.7 *M HClO*_4_ since *P. kluyveri* cells were sticking to the side of the polypropylene tubes during the initial three wash steps with 0.7 *M HClO*_4_.

### Proteomics sample preparation

Cell pellets were prepared and stored as described above. 1 *mL* of lysis buffer (25 *mM* Ammonia Bicarbonate (AmBic) with protease inhibitors (Roche complete protease inhibitor tablets)) was used to resuspend the thawed pellets and the sample was transferred to a 2 *mL* screw cap vial for bead beading in 2 *mL* Precellys vials prefilled with 0.5*mm* glass beads. The sample was centrifuged at 4 ^◦^*C* at 2000 x g for 10 minutes to re-pellet the sample. The supernatant was discarded. The pellet was resuspended in 250 *µL* of lysis buffer. Samples were bead-beat using 15 x 30 seconds beating on high setting with 1 minute cooldown period on ice between each beating cycle. The sample was centrifuged at 4 ^◦^*C* at 13000 x g for 10 minutes to re-pellet the sample. The supernatant was collected and retained. The debris was resuspended using 250 uL of fresh lysis buffer and extracted by piercing a hole at the bottom of the bead-beating tube and centrifuged at 4 ^◦^*C* at 4000 x g for 10 minutes into a collection tube. This was combined with the previously collected supernatant. For each sample, protein concentration was determined using the Bradford Assay. For digestion, from each sample 100 *µg* total protein was taken and diluted to a total volume of 160 *µL* using 25 *mM* AmBic. Proteins were denatured by adding 10 *µL* of 1 % Rapigest solution and heating at 80 ^◦^*C* for 10 min. Cysteine reduction was performed by adding 10 *µL* of an 11.1 *mg* · *mL*^−1^ DTT in 25 *mM* AmBic and incubated at 60 ^◦^*C* for 10 minutes with 450 rpm. Protein alkylation was performed by adding 10 *µL* of a 46.6 *mg* · *mL* iodoacetamide in 25 *mM* AmBic and incubated for 30 minutes in the dark. Iodoacetamide was quenched with 9.4 *µL* of 11.1 *mg* · *mL* DTT in 25 *mM* AmBic. Digestion was carried out by adding 10 *µL* of 0.2 *µg/µL* trypsin, at a final sample:trypsin ratio of 50:1) and incubated at 37 ^◦^*C* 16 hours with 600 rpm chilled to 4 ^◦^*C* until further analysis. Rapigest was inactivated by adding 1 *µL* trifluoroacetic acid ensuring a pH < 2 was achieved. Samples were incubated at 37 ^◦^*C* for 45 minutes at 450 rpm. Samples were then spun in a centrifuge at 13000 x g for 15 minutes at 7 ^◦^*C*. Clarified peptides were recovered and retained for MS analysis.

### LC-MS analysis

Base peak intensity (BPI) target was in the order of 10^6^ − 10^7^ for the timsTOF HT™ mass spectrometer (Bruker). To determine injection volumes of each sample (target of BPI of 10^6^ − 10^7^) survey runs of 15 minutes gradient (30 minutes program) were ran and final sample loading volumes were based on the BPIs observed from the survey runs. A final sample loading volume of 2 *µL* was used for each sample. Injected samples were analysed using a nanoElute 2™ system (Bruker, USA) coupled to a timsTOF HT™ mass spectrometer (Bruker). The sample was loaded onto the trapping column (Thermo Scientific, PepMap100, C18,300 *µm* X 5 mm), using “*µlpickup*” mode, for 4X injection volume + 2 *µl* at a loading pressure of 84.7 bar with 0.1 %(v/v) FA. The sample was resolved on the analytical column (PepSep Twenty-Five Series a C18 150 *µm* x 250 *mm* 1.5 *µm* column) using a gradient of 98 % A (0.1 % Formic Acid in HPLC Water) 2 % B (100 % ACN with 0.1 % Formic Acid) to 65 % A 35 % B over 90 minutes (2-hour method). A data-dependent acquisition with a 10-step parallel accumulation serial fragmentation (PASEF) instrument method acquired between m/z 100-1700 with 1*K*0^−1^ start of 0.6 and end with 1.6*V* · *s* · *cm*^−2^ with a desired target intensity of 20,000 using an intensity threshold of 2500.

### Plate reader cultivation

Cultures were grown on SM media, as described for shake flask cultivations, supplemented with different carbon or nitrogen sources. Cells were grown in a 48-well flat-bottom plate, at 30 ^◦^*C* FLUOstar Microplate Reader (BMG, Germany) with orbital shaking at 700 rpm. Growth was measured by OD600 with 5-minute intervals. Early exponential growing cultures were washed in nitrogen-or carbon source-free SM prior to adding the cultures to the respective media type. Media supplemented with amino acid contained 1.3 *mM* nitrogen and media supplemented with different carbon sources contained 66 *mM* carbon of tested nitrogen or carbon source.

### Cellular volume quantification

To quantify cellular volumes of *P. kluyveri* we used a combination of fluorescent imaging of the cells and cell segmentation [44]. Samples for image analysis were taken from either mid-exponential or steady-state bioreactor cultivations. Cells were incubated with 300 *mM* (*P. kluyveri*) MitoTracker Red CMX-H2Ros (Invitrogen). The MitoTracker concentrations were optimized to fall in between the 16-bit range of the final images. The cells were incubated for 30 minutes with Mitotracker at 30^◦^*C* in destined media type at 200 rpm in the dark. Cells were fixed by incubation with 4 % paraformaldehyde (v/v) for 45 minutes with regular shaking. After fixing, cells were washed twice in cold buffer B (1.2*M* sorbitol in 100*mM KHPO*_4_, pH 7.5), 4000 rpm for 3 minutes at 4 ^◦^*C*. Finally, cells were resuspended in 125 *µl* buffer B or the remainder of the supernatant. Stained, fixated, and washed cells were placed on an 18 mm poly-L-lysine coverslip, in a 12-well plate, for 1h at 4 ^◦^*C*. Subsequently, the coverslip was washed with 2 *mL* buffer B. The cover slip was attached to the object glass after being dipped in 100 % ethanol, air-dried, and inverted cell-side down on a droplet of DAPI-containing mounting solution. After at least 20 hours of incubation, object glasses were sealed using clear nail polish.

Fluorescent images were acquired by a BX63 wide-field epifluorescence microscope (Olympus) equipped with APO-chromatic 100x objective, Orca Fusion camera (Hamamatsu), Tungsten-Halogen lamp, and SOLA FISH LED lamp combined with Olympus Cell Sens Dimension software (Version 2.3 Build18987). Olympus immersion oil was used. The microscope was focused through the DAPI channel (F36-500 DAPI HC Brightline Bandpass Filter), and 80 z-stack images were acquired in the CY3 (F36-542 Cy3 HC BrightLine Filter)(20-200 ms) and DAPI channel (10-20 ms). Exposure times were adjusted in order to capitalize on the full dynamic range of the camera. CellSens Dimension software was used to convert from Olympus .vsi-format to .tif-format. Maximum projection images were created using FIJI (Build c89e8500e4).

A custom data analysis pipeline was created to estimate volumes of cells (see Supplementary Material). Cells were segmented with a custom-trained neural network which used original fluorescent image of mitochondria (CY3-channel). To determine the final cell volumes, the cell masks (from segmentation) were used to count the number of pixels annotated as cell.

### Electron microscopy

Samples for electron microscopy (EM) analysis were acquired from bioreactor cultivations. Directly after sampling, samples were quenched in 2.5 % glutaraldehyde (Merck) in in 0.1 *M* cacodylate (Sigma-Aldrich) buffer (pH 7.4) for an hour at room temperature. This sample was washed twice in Buffer B, and the cell wall was partially digested with lyticase (brand) (25U per OD600) at 37^◦^*C*. Lyticase was removed and the samples were washed twice in Buffer B before being stored in 2.5% glutaraldehyde in 0.1 *M* cacodylate (Sigma-Aldrich) buffer (pH 7.4) at 4^◦^*C* until further processing. Samples were subsequently washed three times with 0.1 *M* cacodylate, pH 7.4 and post-fixed in 1 % *OsO*_4_ (EMS) / 1 % *KRu*(*CN*)_6_ (Sigma Aldrich) for 60 min. Cells were spun down in low-melting point agarose (1 %; Sigma Aldrich) in a swing-out centrifuge at 10000 rpm, 3 min. The pellets were dehydrated by a series of incubations in increasing ethanol concentrations up to 100 % ethanol, followed by rinsing in propylene oxide (EMS) and incubating for 30 minutes in 1:1 mixture of EPON resin:propylene oxide. Pellets were transferred to fresh EPON and moved to embedding molds for resin polymerization for 36 h at 65^◦^*C*. Ultrathin sections were cut at an Ultracut (Reichert-Jung) microtome at 70 − 90 *nm* thickness, and collected on 100 mesh copper EM grids. The sections were contrasted by 4 % *Nd*(*CH*_3_*COO*)_3_ and Reynolds lead citrate in an AC20 (Leica) ultrastainer before imaging on a Tecnai12 (ThermoFisher/FEI) TEM at 100 kV with a Veleta (EMSIS) CCD sidemounted camera at 9900 and 20500 times magnification. Mitochondria were recognized as organelles with a double membrane structure and internal cristae. Mitochondria were quantified by manual segmentation of the mitochondria from the total cell using Napari [45].

### GEM reconstruction

Template-based reconstruction of *P. kluyveri* metabolism was done using MetaDraft (version 0.9.0) [46], with a pipeline previously described in [47]. In short, selected yeast GEMs were ranked in terms of template preference based on MEMOTE score [48] and phylogenetic distance [49]: (1) *Saccharomyces cerevisiae* Yeast8.4.2 [50]; (2) *Issatchenkia orientalis* model iLsor850 [51]; (3) *Kluyveromyces marxianus* model iSM996 [52]; (4) *Yarrowica lipotica* model iYali4 [53]; (5) *Komagataella pastoris* model iMT1026 v3.0 [54]. Thereafter, the draft model was subjected to manual curation. The biomass equation of the model is based on Yeast8.4.2 formulation, with adjustments from biomass composition data from this study.

### pcPichia reconstruction

pcPichia was constructed following the pcYeast8 construction pipeline [13]. For an elaborate description of pcPichia reconstruction and parameterization, see the Supplementary Notes.

### Software

Model construction and optimization were performed with CBMPy package (version 0.8.1) [55] in a Python 3.7.9 environment with the IBM ILOG CPLEX Optimization Studio (version 12.10.0) and SoPlex (version 6.0.2) [56] as the low- and high-precision LP solver, respectively. The dual (-o flag) and primal (-f flag) feasibility tolerance was set to 10^−16^ in SoPlex. R (version 4.1.0) and Python (version 3.7.9) were used for further data analysis and visualization.

## Acknowledgements

This research was supported by the Dutch Research Council grant no. ENPPS.LIFT.019.005, a public private-partnership grant supported by Novonesis. We thank SURF (www.surf.nl) for the support in using the National Supercomputer Snellius. We thank Jack Pronk for the fruitful discussions. We thank Marcel Vieira Lara and Marieke Warmerdam for helping with biomass composition measurements. We thank Christiaan Mooijman for the valuable discussions on bioreactor cultivations. We thank Sander Otterdijk for his valuable input on cell segmentation and Kelly van Rossum for her input on fixation protocols. We thank J.R.T. van Weering en M.P. Dekker from EM facility Amsterdam for their work on electron microscopy. We thank P. Brownridge and C. Kennedy from the Center of Proteome Research in Liverpool for their work on proteomics.

## Author contributions statement

Conceptualization, funding acquisition and supervision: B.T., F.J.B., J.K.A.; experimental data collection: J.B., M.H., G.A.M., T.D.V., K.S.; experimental data analysis: J.B., M.H., G.A.M., T.D.V., K.S.; computational modeling: J.B., P.G.; formal analysis: J.B., P.G., J.vH., F.J.B., B.T.; writing – original draft: J.B.; writing – editing: J.B., P.G., B.T. All authors have read and approved the manuscript.

## Supplementary material

Supplementary Data 1 – Physiology

Supplementary Data 2 – GEM

Supplementary Data 3 - Proteome

Supplementary Data 4 - Cellular and mitochondrial quantification

Supplementary Data 5 - pc-model

## Data and code availability

Genome: will be made publicly available when published Models, scripts, and genome annotation: will be made publicly available when published

## Competing Interests

The authors declare no competing interests.

## Notes

### Competing Interest Statement

The authors have declared no competing interest.

