## Supplementary Material for "Mitochondrial efficiency determines Crabtree effect across yeasts"

Julius Battjes *et al.*

### 1 Supplementary Figures

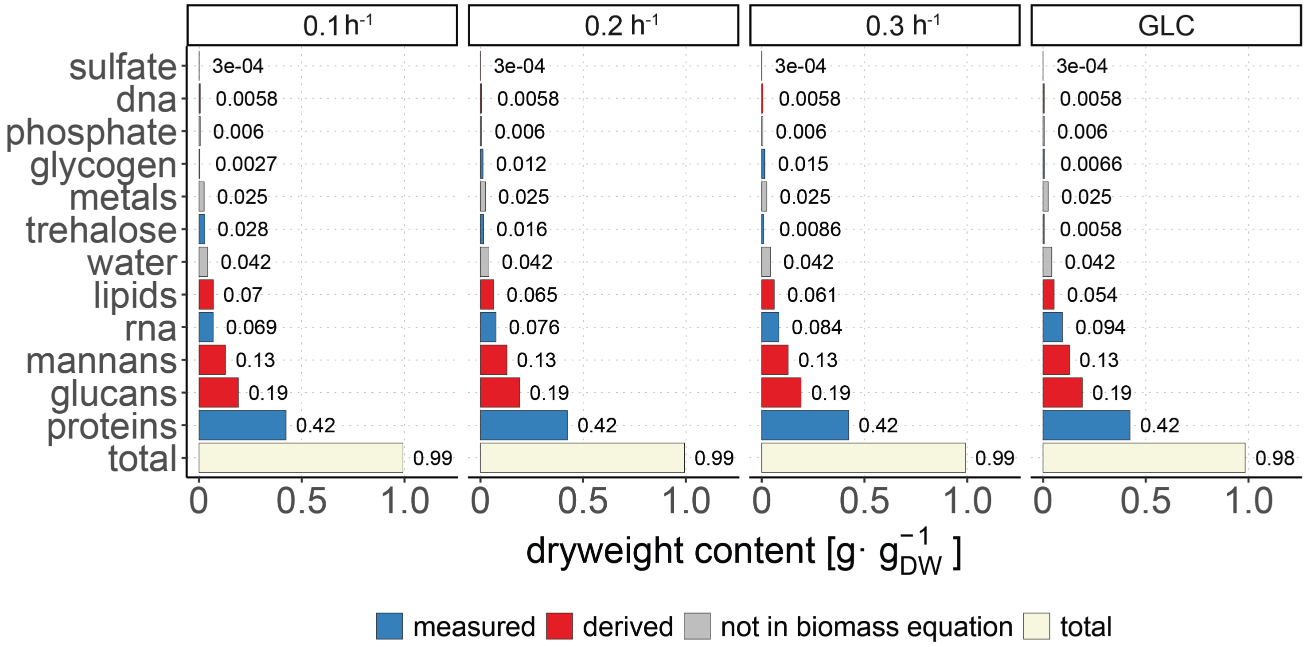


**Supplementary Figure 1.** Total biomass composition as determined per condition for *P. kluyveri*. Amounts of components that were not determined experimentally were inferred from literature values (Canelas et al., 2011). The derived biomass equation for *P. kluyveri* in glucose batch was used as for the biomass equation of its GEM (see Supplementary Notes for details).

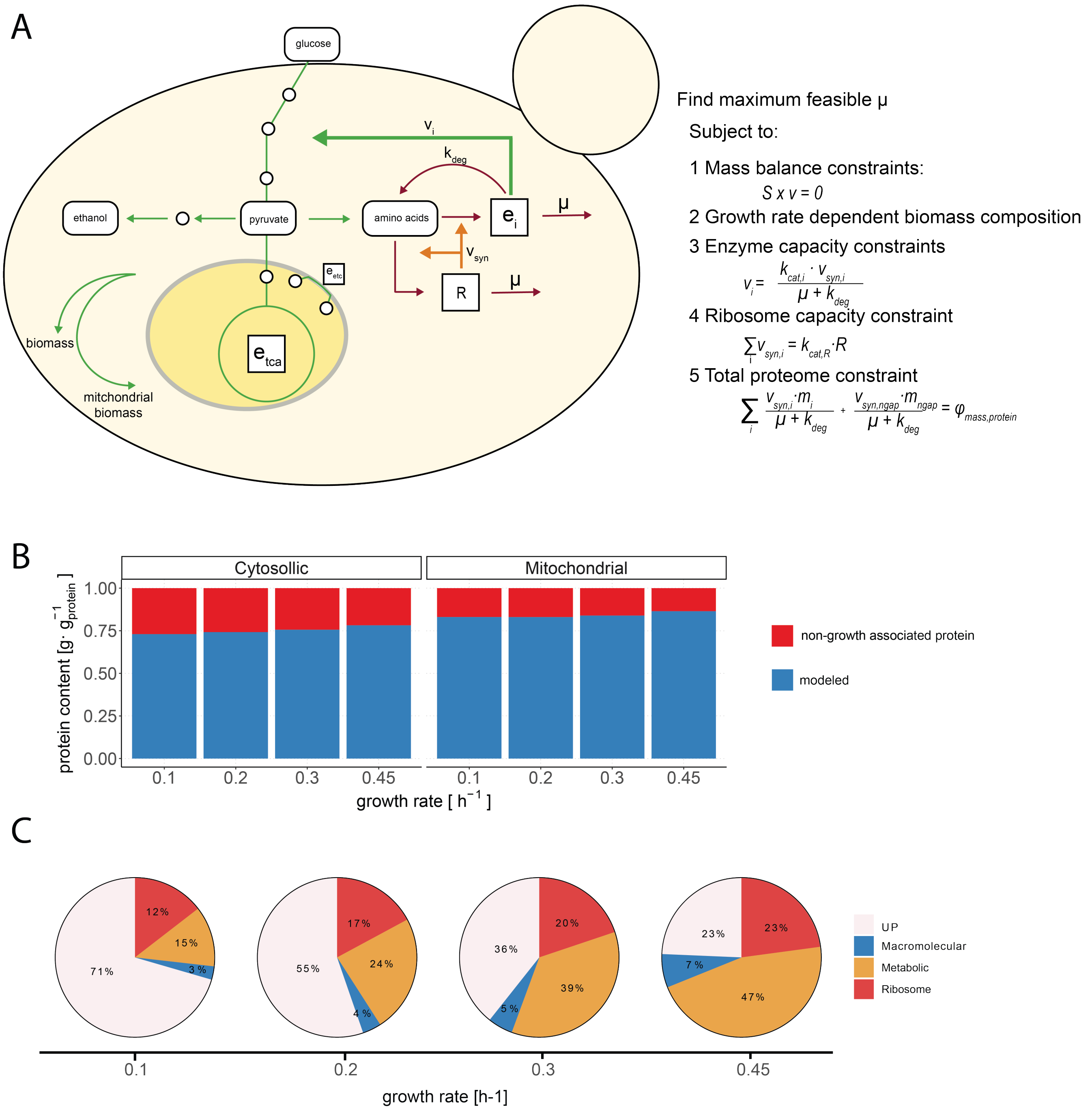


**Supplementary Figure 2.** The model structure and predicted proteome partitioning in the proteome-constrained model of *P. kluyveri*, pcPichia. (A) schematic overview of the pc-model framework used in this study, adapted from Elsemman et al. (Elsemman et al., 2022; Grigaitis et al., 2023). Metabolites (circles) are made in the stoichiometric network from external nutrients, which are used to produce biomass. (B) The amount of non-growth associated protein (NGAP) was derived by comparing proteome dataset to proteins in the pc-model of *P. kluyveri*. Minimal amount of NGAP in both the cytosol (22%) and mitochondria (14%) were found at $\mu_{max}$. (C) The predicted proteome partitioning at increasing growth rates. The predicted fraction of non-growth-associated protein (NGAP), a proxy for protein which is not accounted for with specific reactions in the pcPichia model, decreases with increasing predicted growth rate until it reaches the predetermined minimum fraction ($NGAP_{min}$). Fractions of metabolic protein, ribosomes, and macromolecular protein (protein turnover related), increase with increasing growth rate.

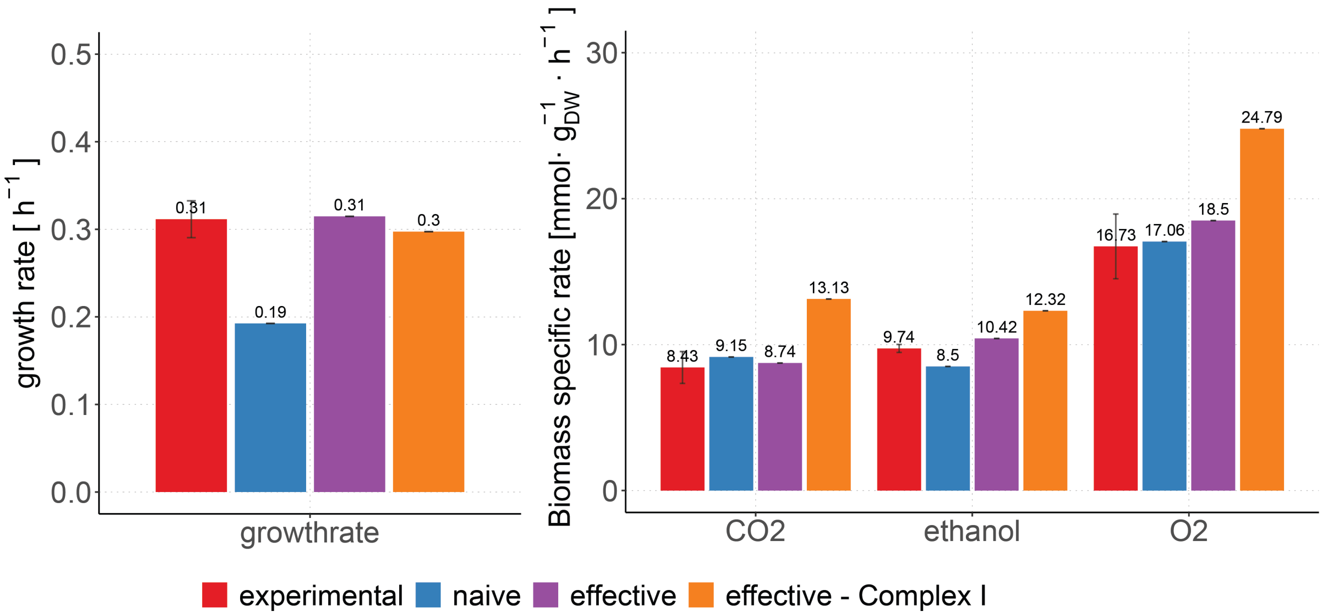


**Supplementary Figure 3.** pcPichia predictions compared to experimental growth of *P. kluyveri* in ethanol batch conditions. We corrected $NGAP_{min}$ for pcPichia (effective) from 0.19 $g \cdot g_{protein}^{-1}$ to 0.30 $g \cdot g_{protein}^{-1}$ to match the maximum observed growth rate, and applied this $NGAP_{min}$ to all ethanol grown models (right panel).

#### 2 Supplementary Tables

**Supplementary Table 1.** Observed relative mitochondrial sizes of *S. cerevisiae* from literature. NA, not available.

| carbon source | media | method | $\psi_{mitochondria}$ | n | growth rate $[h^{-1}]$ | reference |
|---|---|---|---|---|---|---|
| glucose | complex | Su9-mCherry | 0.025 - 0.05 | >27 | NA | (Tsuboi et al., 2020) |
| glucose | synthetic | CSLM | 0.074 | 9-13 | NA | (Visser et al., 1995) |
| ethanol | synthetic | CSLM | 0.063 | 10-11 | NA | (Visser et al., 1995) |
| glucose | synthetic | pMitoLoc | 0.0477 | 50 | NA | (Bartolomeo et al., 2020) |
| ethanol | synthetic | pMitoLoc | 0.35 | 50 | NA | (Bartolomeo et al., 2020) |
| glucose/ethanol | synthetic | pMitoLoc | 0.1116 | 50 | NA | (Bartolomeo et al., 2020) |
| lactate | complex | EM | 0.12 | 235 | 0.3 | (Grimes et al., 1974) |

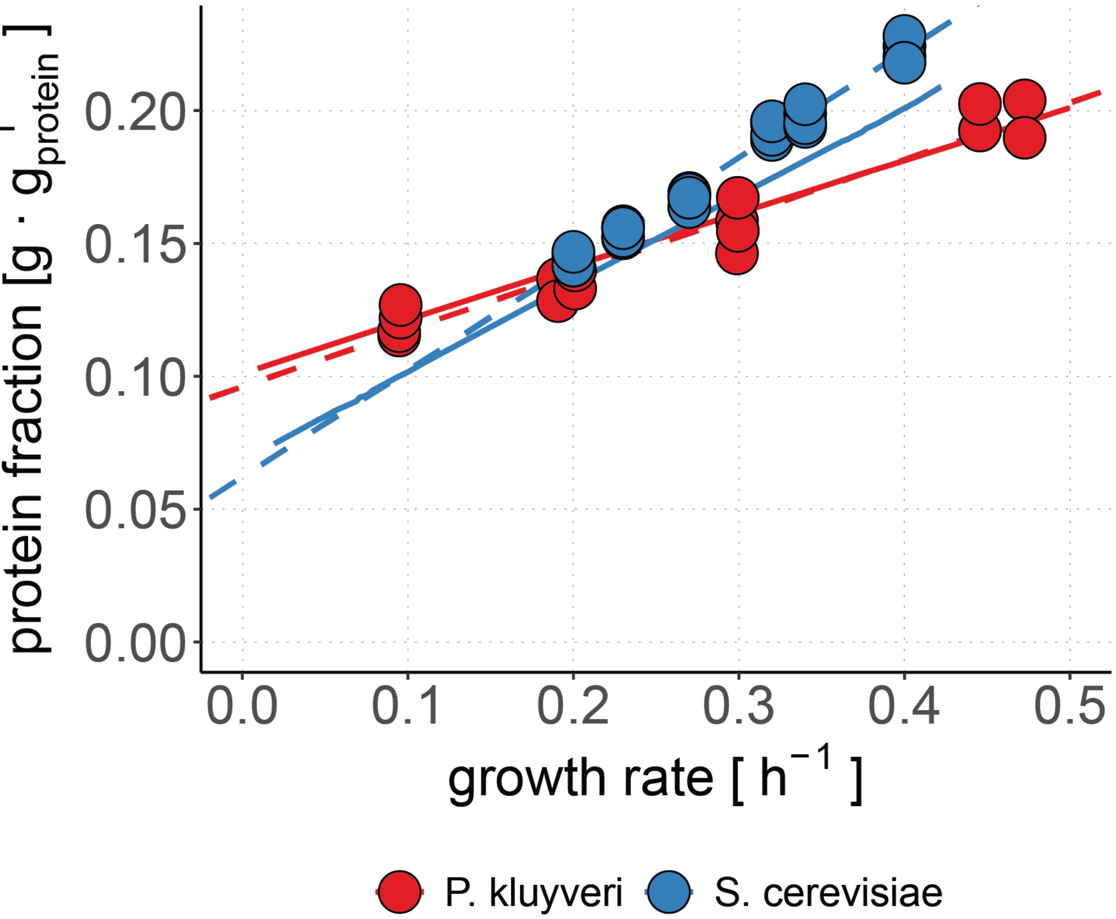


**Supplementary Figure 4.** Ribosomal proteome fraction of *P. kluyveri* and *S. cerevisiae* compared to pcPichia and pcYeast. For pcPichia, ribosomal elongation rate (16 $aa \cdot s^{-1}$) is fitted (solid line) to experimental data (circles). For pcYeast, ribosomal elongation rate (10.5 $aa \cdot s^{-1}$) is derived from literature (Elsemman et al., 2022). Linear regression fit of the experimental data is visualized with dashed lines.

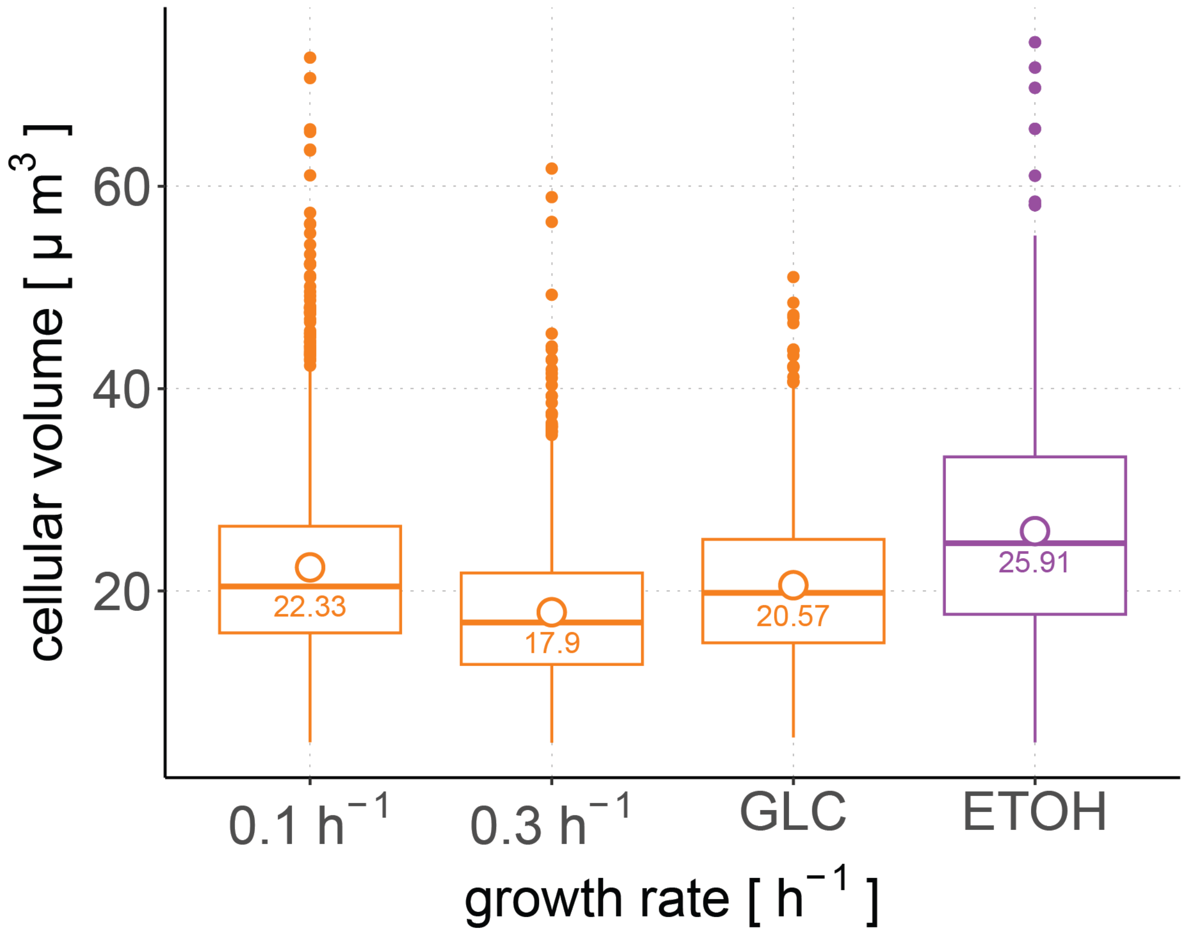


**Supplementary Figure 5.** Cellular volumes of *P. kluyveri* cultured in aerobic, glucose limited ($0.1h^{-1}$ & $0.3h^{-1}$), glucose batch (GLC), and ethanol batch (ETOH) conditions. Mean values are displayed per condition by number and open dot. 1200 cells were used per analysis for $0.1h^{-1}$, 1609 for $0.3h^{-1}$, 1620 for glucose batch and 2494 for ethanol batch conditions.

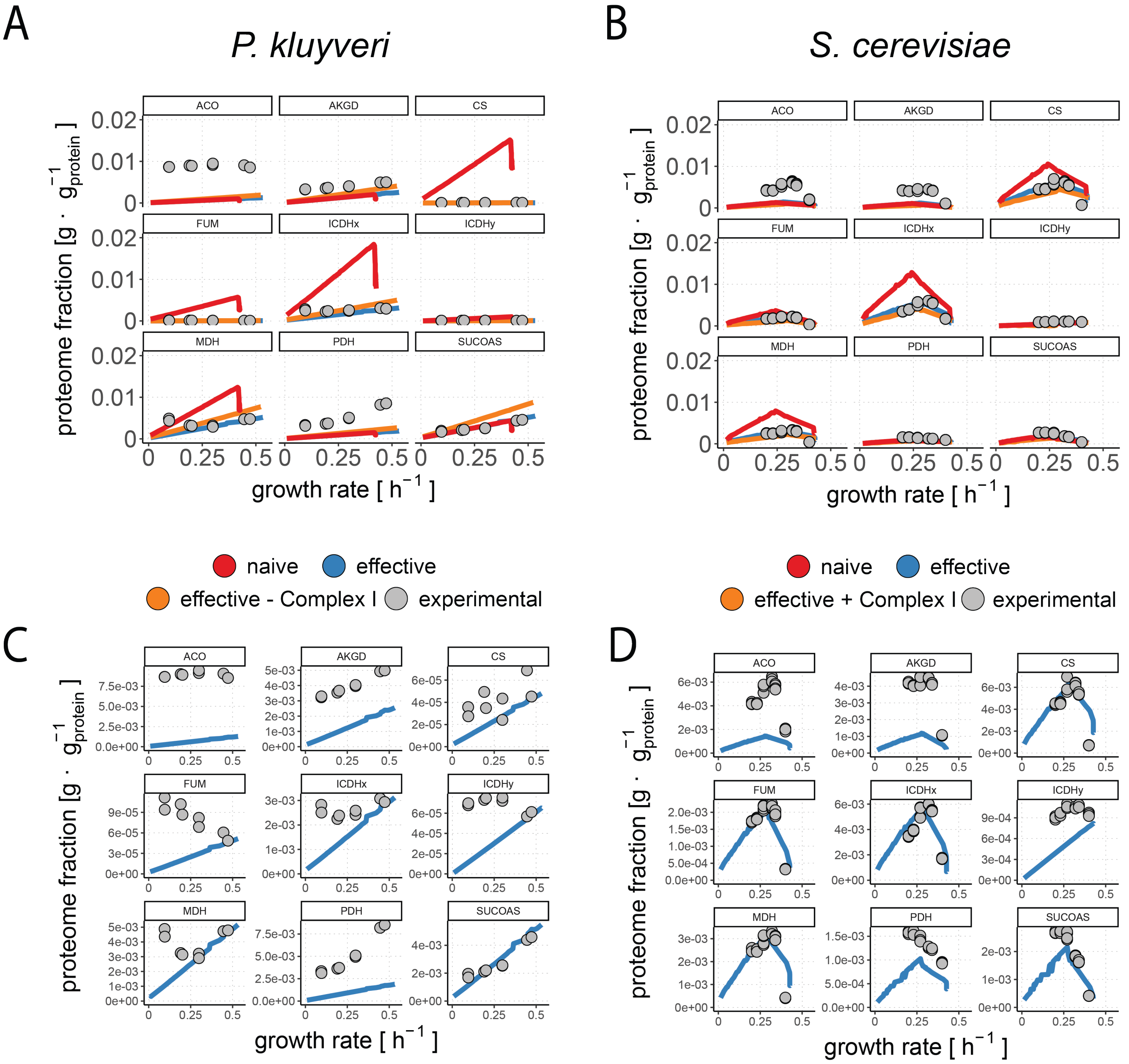


**Supplementary Figure 6.** Predicted and measured levels of TCA cycle proteins. (A) Comparison of protein fractions between different pcPichia models (lines) and experimental data (dots). (B) Comparison of protein fractions between pcYeast configurations (lines) and proteome data (dots) from Elsemman et al. (Elsemman et al., 2022). (C) Same as in A but enlarged for pcPichia (effective). (D) Same as in B but enlarged for pcYeast (effective).

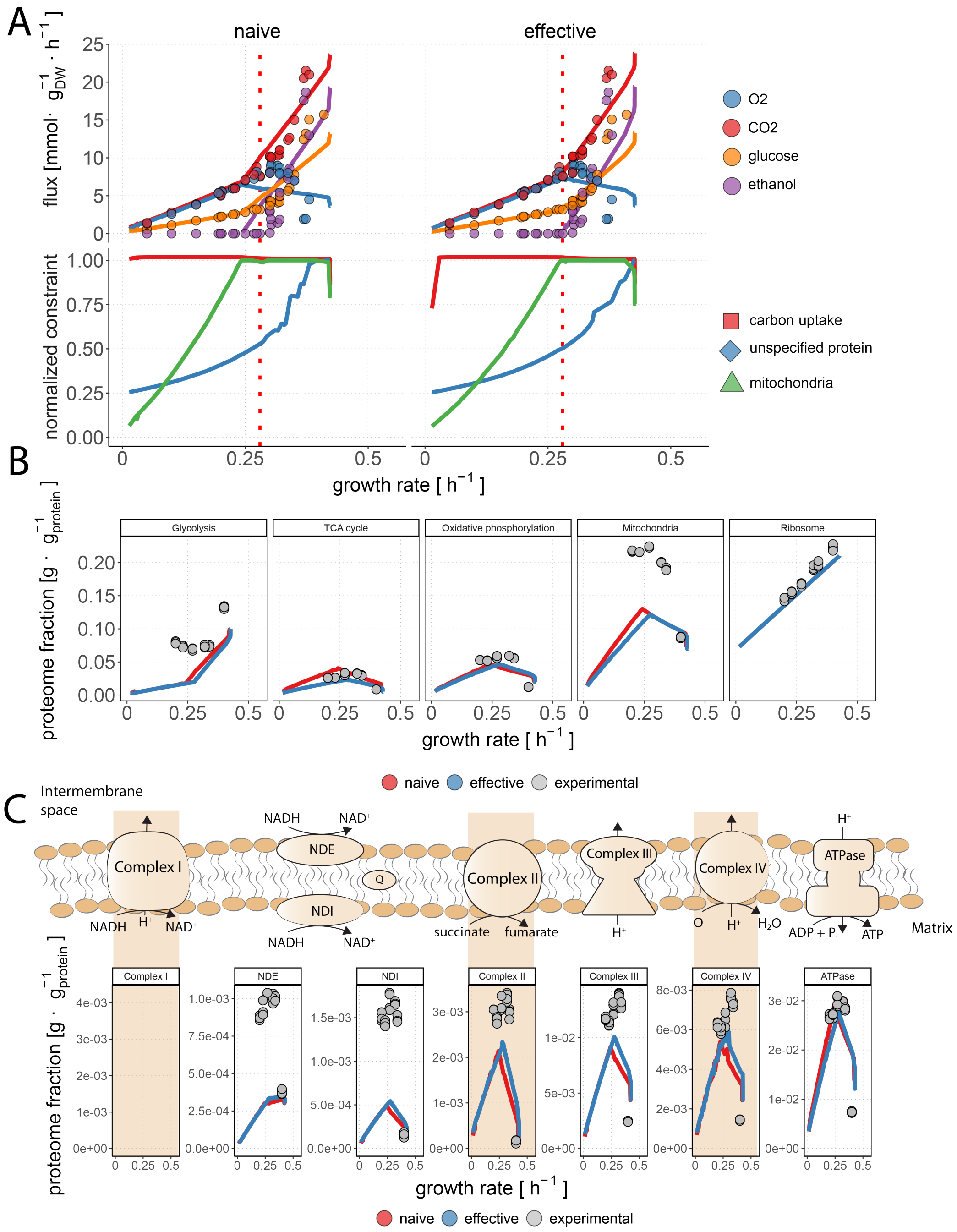


**Supplementary Figure 7.** pcYeast favours fermentation over respiration regardless of Complex I presence. (A) (top) Flux profiles of pcYeast (lines) and experimental *S. cerevisiae* data of glucose limited chemostat and glucose batch experiments (points) (Elsemman et al., 2022; van Hoek et al., 2000). In pcYeast (effective), catalytic constants were changed by comparison between pc-model predictions and measured proteome (Table 3). (bottom) Constraint evaluation of pcYeast for total proteome, plasma membrane, and mitochondrial volume constraints. When the capacity of the compartment is exceeded, the expression of the constraint equals one, and therefore the compartment capacity limits growth. (B) Pathway expression of pcYeast predictions compared to measured proteome fractions. (C) Expression levels of individual oxidative phosphorylation proteins for the different pcYeast configurations.

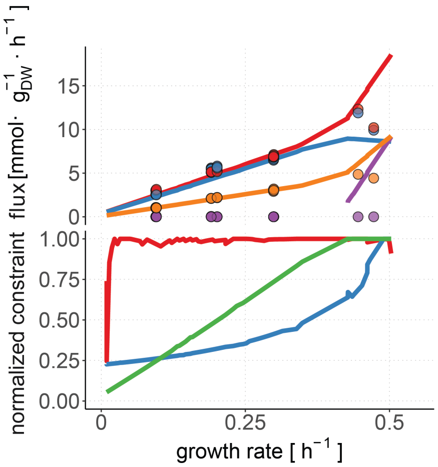


**Supplementary Figure 8.** Uncorrected mitochondrial volume constraint results in limited mitochondrial volume for pcPichia. (top) Flux profiles of pcPichia (lines) and experimental *P. kluyveri* data of glucose limited chemostat and glucose batch experiments (points). Here, we show data of pcPichia (effective) with a mitochondrial volume fraction uncorrected with respect to relative mitochondrial volume differences between *P. kluyveri* and *S. cerevisiae* (See Supplementary Notes for details). (bottom) Constraint evaluation of pcPichia for total proteome, plasma membrane, and mitochondrial volume constraints. The expression of the constraint equals one when the maximal capacity of the compartment is reached, and therefore the compartment capacity limits growth. Visible here for first mitochondrial volume constraint followed by the total proteome constraint.

#### 3 Supplementary Notes

In this study we constructed a genome scale metabolic model (GEM) of *P. kluyveri*. We manually curated the metabolic network of *P. kluyveri*, and derived a biomass equation using our original experimental data. We have used the GEM to predict the C- and N-sources that *P. kluyveri* can use for growth, and predict flux profiles under different nutrient-limited conditions. To extend our analysis, and probe the resource allocation aspect, we also constructed a proteome-constrained model (pc-model), an extension of a conventional GEM that introduces (i) couples metabolic fluxes with enzyme demands, and (ii) adds additional set of constraints describing limited capacity of storing proteins in cell compartments. In these Supplementary Notes we will explain how we constructed the GEM, followed by an overview of the pc-model.

##### 3.1 *P. kluyveri* GEM construction

A GEM is both a tool to study physiology and a repository of knowledge (Somerville et al., 2022). In this study, we constructed the first GEM of *P. kluyveri* from its sequenced genome. First, we performed a homology search between the annotated genome of *P. kluyveri* and genomes of yeast species *S. cerevisiae* (Lu et al., 2019), *Issatchenkia orientalis* (Suthers et al., 2020), *Kluyveromyces marxianus* (Marcišauskas et al., 2019), *Yarrowica lipotica* (Kerkhoven et al., 2016), and *Komagataella pastoris* (Tomàs-Gamisans et al., 2017) to map orthologous genes to gene-reaction associations in their respective GEMs, and so to obtain a draft GEM (Olivier, 2018). The *Saccharomyces cerevisiae* GEM Yeast8.4.2 (Lu et al., 2019), was used as the primary source of reactions and the basis of the biomass equation.

*P. kluyveri* grows on minimal media with a single known vitamin auxotrophy for pyridoxine. Thus, in modeling terms, the GEM of *P. kluyveri* should be able to produce every precursor for biomass production except pyridoxine (Fig. 1). With that in mind, we have curated the draft GEM till the GEM could produce biomass from minimal media with vitamin supplementation.

An accurate biomass equation and ATP maintenance parameters are essential for good quantitative agreement of model predictions to experimental data. We formulated a biomass equation of *P. kluyveri* on the basis of Yeast8.4.2 biomass equation, using our own measurements of protein, RNA, trehalose, and glycogen content (see Section Section 3.2.1) (Lu et al., 2019).

We also included Complex I in *P. kluyveri* GEM (Fig. 2 B). A stoichiometry of 4 $H^{+}/2\ e^{-}$ exported from the mitochondrial matrix was set for Complex I. This stoichiometry has been reported for bacteria, mammals, and yeast (Kaila and Wikström, 2021). The stoichiometry of mitochondrial ATPase was set to 4 $H^{+}$ imported into the mitochondria per ATP (Petersen et al., 2012). The P/O ratio increases from 0.95 $mol_{ATP} \cdot mol_{oxygen}^{-1}$ (using NDI) to 1.84 $mol_{ATP} \cdot mol_{oxygen}^{-1}$ when electron transport chain complex I is used. This P/O ratio for *P. kluyveri* GEM with NDI is equivalent to the P/O ratio of 0.95 of *S. cerevisiae* (Verduyn et al., 1990).

We next fitted the maintenance parameters for the *P. kluyveri* GEM using the data on glucose-limited and -excess growth conditions (Fig. 1). We determined the non growth-associated ATP maintenance (NGAM) of 3 $mmol_{ATP} \cdot g_{dw}^{-1} \cdot h^{-1}$ and growth-associated maintenance (GAM) of 50 $mmol_{ATP} \cdot g_{dw}^{-1}$. In comparison, the NGAM in Yeast8.4.2 was 0.7 $mmol_{ATP} \cdot g_{dw}^{-1} \cdot h^{-1}$ and GAM of 55.3 $mmol_{ATP} \cdot g_{dw}^{-1}$. The difference is more evident when computing the factual requirements (sum ATP hydrolysis flux through NGAM and GAM) at a fixed growth rate. For instance, at a growth rate of 0.1 $h^{-1}$, where both yeast are completely respiratory, this would result in a total ATP maintenance requirement of 8 $mmol_{ATP} \cdot g_{dw}^{-1}$ and 6.05 $mmol_{ATP} \cdot g_{dw}^{-1}$ for *P. kluyveri* and *S. cerevisiae*, respectively. *P. kluyveri* and *S. cerevisiae* have similar flux profile in the respiratory regime of *S. cerevisiae* (Fig. 1 G) despite that it is likely that *P. kluyveri* has an higher $Y_{ATP/glucose}^{theoretical}$ due to additional proton-motive force generated by respiratory Complex I usage.

The curated GEM was then compared to experimental data acquired in this study. The curated GEM correctly predicted the growth/no-growth phenotype in 86% of the measured carbon and nitrogen sources (Fig. 1 C) and quantitatively predicted major metabolic fluxes in aerobic growth on glucose minimal media (Fig. 1 F).

#### 3.2 *P. kluyveri* pc-model construction

For analysing which proteome constraints govern metabolism in *P. kluyveri*, we constructed a pc-model from its GEM. By adding experimentally determined cellular constraints into the growth maximization problem, we can see how the cell steers its investments of limited resources (e.g. biomass precursors, energy, and cellular space) to maximize its growth rate. pcPichia was reconstructed with the published pipeline for the pcYeast model (Grigaitis et al., 2023).

To construct a pc-model from a GEM, we have to add several additional layers of information, such as descriptions of protein composition and turnover, as well as constraints reflecting limited protein capacity of compartments (Supplementary Figure 2 A). Enzyme capacity constraints relate the catalytic rate constant ($k_{cat}$) of an enzyme to the flux, i.e. how much enzyme is required ($v_{syn,i}$) to run a reaction. Enzymes are diluted by growth ($\mu$) and part of the enzymes are degraded ($k_{deg}$). This enzyme synthesis flux ($v_{syn,i}$) is coupled to the ribosome ($R$) capacity constraint. Here, the amount of ribosome required for protein translation ($\sum_i v_{syn}$) is related to the $k_{cat,R}$ of the ribosomes. At last, the cell has a finite protein content, defined by the total proteome constraint. The total proteome consists of two protein pools, the modelled enzymes specified in the pc-model ($\sum_i v_{syn}$) and the non-growth associated protein ($\sum_i v_{syn,ngap}$), which are constrained by the total amount of protein within the cell ($\phi_{mass,protein}$).

For a more comprehensive explanation we refer to Elsemman et al. (Elsemman et al., 2022), and here we cover the changes implemented in pcPichia to reflect aspects specific to *P. kluyveri* and how they were implemented in the model.

##### 3.2.1 *Condition-dependent biomass composition*

We measured the biomass composition of *P. kluyveri* in the glucose-limited and excess conditions, and we observed that biomass content of some precursors varies with growth rate in *P. kluyveri* (Supplementary Figure 1). For *P. kluyveri* GEM we used the biomass composition determined in glucose batch (Supplementary Figure 1). In the pc-model, the growth rate is a parameter in the linear problems, therefore we can (and should) adapt the biomass equation for specific growth rate aiming for a better quantitative agreement of predictions with experimental data.

We obtained the following relationships between content and growth rate of protein (Eq. 1), RNA (Eq. 2), trehalose (Eq. 3), and glycogen (Eq. 4) are also shown in (Fig. 1 E).

$$\phi_{mass,protein} = \mu \cdot 0.036 + 0.413 \quad (1)$$

$$\phi_{mass,RNA} = \mu \cdot 0.072 + 0.062 \quad (2)$$

$$\phi_{mass,trehalose} = 0.044 - 0.184 \cdot \mu + 0.220 \cdot \mu^2 \quad (3)$$

$$\phi_{mass,glycogen} = -0.014 + 0.201 \cdot \mu - 0.345 \cdot \mu^2 \quad (4)$$

As protein content only differs slightly with growth rate, an average protein content of 0.423 $g_{protein} \cdot g_{dw}^{-1}$ was used in pcPichia (Supplementary Figure 1). *P. kluyveri* has a lower and growth rate-independent protein content in the biomass, compared to *S. cerevisiae*, where protein content linearly increases as a function of growth rate between 0.35 $g_{protein} \cdot g_{dw}^{-1}$ and 0.53 $g_{protein} \cdot g_{dw}^{-1}$, between $\mu = 0.025\ h^{-1}$ and $\mu \to 0.4\ h^{-1}$ (Canelas et al., 2011) (Table 2).

Biomass content of inorganic ions (metals), and water were not specifically included in the pc-model biomass equation for simplification. We reasoned that incorporation would only require additional transport reactions carrying relatively small fluxes (Canelas et al., 2011). Lipid composition of cell membrane and mitochondrial membrane in pcPichia was adjusted from the pcYeast formulation. When a specific lipid could not be produced by pcPichia, it was removed from the equation. The amount of each lipid in the biomass was proportionally adjusted arrive to the amounts consistent with the composition of 1 $g_{lipid}$. Amounts of mannans and glucans were derived from measurements of *S. cerevisiae* and incorporated in a ratio of 1:1.5, respectively, similar to *S. cerevisiae* (Canelas et al., 2011). Altogether, with the relationships implemented, the mass fraction, explicitly represented by the biomass equation is stable with varying growth rate (Supplementary Figure 1).

###### 3.2.2 ATP maintenance

pcPichia already accounts for a part of the ATP maintenance by explicitly defining ATP costs of protein translation, folding, and degradation. The maintenance coefficient of pcPichia was calibrated to correct for explicitly-defined ATP requirements, using the data of *P. kluyveri* growth in glucose-limited and -excess conditions, same as described above for the GEM. We titrated the parameters in the version of pcPichia model with two energy generation-relevant considerations: (i) we used corrected mitochondrial volume constraint of 0.791 $\mu m^3$ (Table 2), (ii) the mitochondria harbored proton-pumping respiratory Complex I, and (iii) effective $k_{cat}$'s were used (Table 3). This led to a GAM estimate of 40 $mmol \cdot g_{dw}^{-1}$ and an NGAM of 0.5 $mmol \cdot g_{dw}^{-1} \cdot h^{-1}$. For *S. cerevisiae*, an GAM of 24 $mmol \cdot g_{dw}^{-1}$ and an NGAM of 0.168 $mmol \cdot g_{dw}^{-1} \cdot h^{-1}$ were used in (Grigaitis et al., 2023). This big difference in maintenance coefficients accounts for the higher P/O ratio of pcPichia (1.84 $ATP \cdot O^{-1}$) compared to pcYeast (0.95 $ATP \cdot O^{-1}$), while the observed fluxes (and yields Yx/s?) are similar for both organisms in respiratory regime ($\mu < \mu_{critical}$ for *S. cerevisiae*).

##### 3.2.3 Non-growth associated protein

The pc-model accounts only for known metabolic (and protein turnover-related) proteins in the proteome of *P. kluyveri*. In addition, the pc-model predicts the minimum amount of protein required for each reaction, running at $v_{max}$. There thus remains a fraction protein that does not actively contribute to growth: proteins performing other functions, such as signaling, structural; or metabolic proteins expressed but not used at their full flux capacity (observed $v << v_{max}$).We observed that the measured protein content in biomass remains stable for *P. kluyveri* at different growth rates (Eq. 1), and this means that the rest of the proteome space that has to be filled with protein to maintain the stable total protein content in biomass. We have thus defined an artificial non-growth associated protein (NGAP) to represent proteins without metabolic function and/or proteins that are undersaturated. The NGAP expression typically decreases with increasing growth rate and reaches a pre-defined minimal level (Supplementary Figure 2 B): then we render that the proteome capacity constraint actively limits growth. This means that all available proteome space is used for growth, and any change in protein expression must come at the cost of another protein.

Here, we used the same definition for NGAP in pcPichia as for pcYeast (Elsemman et al., 2022). This composition of NGAP resembles an average protein with a total mass of 39.8 kDa and chain length of 450 amino acids. The minimal amount of NGAP in the cell was set to the amount of protein not represented by the model. This number was derived by computing the fraction of the proteome that is not represented within pcPichia. This resulted in a minimum of 22 % of NGAP at $\mu_{max}$ (Supplementary Figure 2 C). The mitochondria have their own modelled fraction of NGAP, which was found to be 14% at $\mu_{max}$ (Supplementary Figure 2 C). Compared to *S. cerevisiae*, these numbers are fairly similar (Table 2).

#### 3.3 Enzyme and pathway curation

The enzyme capacity constraint states that a reaction can only carry flux if respective enzymes are synthesized. The required enzyme abundance to sustain a flux depends on the catalytic first order rate constant ($k_{cat}[h^{-1}]$) (Eq. 5).

$$v_i = \frac{k_{cat,i} \cdot v_{syn,i} \cdot f}{\mu + k_{deg}} \tag{5}$$

In which $v_i$ is the flux through enzyme i, $v_{syn,i}$ is the protein synthesis flux, and $k_{deg}$ is the degradation rate of enzymes. We set $k_{deg}$ to $0.043h^{-1}$, which is the degradation rate determined for *S. cerevisiae* at $\mu = 0.1h^{-1}$ (Lahtvee et al., 2017). To calculate the minimal protein requirement, i.e. under assumption that each enzyme operates at its $v_{max}$, enzyme saturation function f is set to 1.

Generally, we used the $k_{cat}$ values of pcYeast for their respective orthologues in pcPichia. However, special attention was given to the annotation and assignment of $k_{cat}$ for specific pathways, described here.

##### 3.3.1 Oxidative phosphorylation

Oxidative phosphorylation of *P. kluyveri* was subjected to additional manual curation. To map the subunits of respiratory Complex I, *P. kluyveri* proteome sequences were compared to protein sequences of *Yarrowia lipolytica* using BLAST and Inparanoid, for which sequences were derived from Uniprot (Bateman et al., 2020; O'Brien, 2004). Out of the 14 proposed

Complex I core subunits, two could not be identified in the *P. kluyveri* proteome (NULM and NU2M). As for the accessory subunits, each of the 23 subunits had at least one positive hit. For modelling purposes, the median size of 1.067 MDa of all complex I sizes was picked for the pcPichia model, close to the proposed size of $\tilde{1}$ MDa (Sharma et al., 2019). Initially, we set the $k_{cat}$ of 370 $s^{-1}$ for Complex I, measured for *Komagataella pastoris* (Bridges et al., 2009). Later in the process, the $k_{cat}$ of Complex I was adjusted to 720 $s^{-1}$ to fit the observed Complex I expression levels at $\mu_{max}$ (Fig. 3 C).

For the rest of ETC complexes, the subunit stoichiometry was taken from pcYeast. Complex III, Complex IV and ATPase were manually curated by comparison to *S. cerevisiae* protein sequences. *P. kluyveri* Complex III (227 kDa) and Complex IV (171 kDa) are of similar size to *S. cerevisiae* counterparts (250 kDa and 200 kDa, respectively) in (Schagger, 2000). We adapted $k_{cat}$ from *S. cerevisiae* for complex III and complex IV: 220 $s^{-1}$ and 693.5 $s^{-1}$, respectively, compared to 120 $s^{-1}$ for both enzymes in pcYeast (Grigaitis et al., 2023), these new values were taken from Nisson et al. (Nilsson and Nielsen, 2016). The predicted ATPase size of *P. kluyveri* is 492 kDa, while the ATPase of *S. cerevisiae* is approximately 570 kDa (Velours and Arselin, 2000). This size difference is mainly due to the missing OLI1 (ATP9) gene in the annotated *P. kluyveri* genome. The smallest available size for ATPase in pcPichia is 504 kDa. The size for ATPase for pcYeast was previously set to 326 kDa, this was now changed to 485 kDa. In addition, for pcPichia the $k_{cat}$ of ATPase was increased from the reported 120 $s^{-1}$ to 300 $s^{-1}$, so model predictions at $\mu_{max}$ matched protein expression levels (Förster et al., 2010).

###### 3.3.2 TCA cycle

We also checked if catalytic constants of pcYeast TCA cycle protein were applicable to *P. kluyveri*. One catalytic constant stood out when comparing pc-model predictions with protein expression data (Supplementary Figure 6 A). For citrate synthase (CS) predictions of pcPichia were well above actual protein levels around $\mu_{max}$ (pcPichia naïve). Therefore we changed the catalytic constant of CS from the TCA cycle from 11.2 $s^{-1}$ to 141.3 $s^{-1}$ (latter taken from (Nilsson and Nielsen, 2016)). With the changes implemented, the total proteome fraction attributed to the TCA cycle for *P. kluyveri* drop to near experimental values at $\mu_{max}$ in pcPichia (effective)(Fig. 3 B).

###### 3.3.3 Ribosomes and macromolecular complexes

To assign the correct proteins to the ribosomes and macromolecular complexes in the model, we matched *P. kluyveri* protein sequences with the protein sequences of *S. cerevisiae* using Inparanoid orthology finder (O'Brien, 2004). Ribosomal content of *P. kluyveri* increases with growth rate, according to a linear relationship (Eq. 6):

$$\phi_{ribosome} = \mu * 0.214 + 0.096 \tag{6}$$

The ribosomal protein fraction ($\phi_{ribosome}$) expressed at zero growth, is about 9.6 % of the total expressed protein. The ribosomal elongation rate and inactive fraction were calibrated with pcPichia to resemble the observed ribosomal protein fraction. This resulted in a ribosomal elongation rate ($k_{elongation}$) of 16 $aa \cdot s^{-1}$ and an ribosomal inactive fraction ($\phi^0_{ribosome}$) of 0.1 (Supplementary Figure 4). Compared to *S. cerevisiae*, the elongation rate is lower while the void fraction is higher.

###### 3.3.4 Glucose transport

The glucose concentration in broth from each of the three chemostat cultivations was below 0.1 $mM$, and thus this could have implications to the efficacy of glucose uptake by the cells. We can evaluate the transport affinity using the Monod equation (Eq. 7), i.e. calculate the maximum Monod constant (glucose affinity $K_M$) of *P. kluyveri*.

$$\mu = \mu_{max} \cdot \frac{C_{s,out}}{C_{s,out} + K_M} \tag{7}$$

The highest dilution rate in the chemostat was about 0.3 $h^{-1}$ ($\mu$), while the maximum measured growth rate $\mu_{max} = 0.45 h^{-1}$. We did not measure any residual glucose in the chemostat broth at 0.3 $h^{-1}$ or lower, therefore we take the lower limit of the enzymatic glucose determination assay of 100 $\mu M$ ($c_{s,out}$). This results in a maximum $K_M$ of 50 $\mu M$. Glucose transport of Crabtree negative yeast more often have low Monod constants in a similar range (Urk et al., 1989).

The mode of glucose transport in *P. kluyveri*, i.e. whether proton symport or facilitated diffusion for glucose transport is used, is not known. Moreover, the mode might differ between lower growth rates or $\mu_{max}$ (Urk et al., 1989). Three genes were annotated as glucose transporters in*P. kluyveri* GEM, 2521_g, 1619_g, and 4948_g. It is assumed that at $\mu_{max}$, when glucose is available in ample amounts, facilitated diffusion is the preferred mode of transport. As for the $k_{cat}$ for glucose transport, a general rule of thumb is that the higher the affinity, the lower the $k_{cat}$ (Bosdriesz et al., 2018). The first-order rate constant of the glucose transporter of *S. cerevisiae* in pcYeast is set to 200 $s^{-1}$. As there is no further information on the $k_{cat}$ of glucose transport in *P. kluyveri* is available, the $k_{cat}$ was kept at 200 $s^{-1}$. This $k_{cat}$ both resembles the $k_{cat}$ used in pcYeast and is close to the lower $k_{cat}$ values found for glucose facilitated diffusion in *S. cerevisiae* (Bosdriesz et al., 2018; Grigaitis et al., 2023).

##### 3.4 Formulation of proteome constraints

In pc-models we can put a upper constraint on different protein pools. These constraints can be imposed on the complete proteome and on different organelles. Previously, such a mitochondrial volume constraint showed that *S. cerevisiae* was limited in its mitochondrial synthesis capacity (Elsemman et al., 2022). Therefore, we calculated a mitochondrial volumetric constraint in similar fashion. This volumetric constraint is depended on cell volume, cell density, and mitochondrial protein content. Additionally, cell volume is used to determine the cell membrane space available for transport protein.

###### 3.4.1 Total proteome constraint

To limit total protein content of the cell we impose a total proteome constraint, in which all the produced protein must be equal to the measured protein mass fraction of 0.42 $g_{protein} \cdot g_{dw}^{-1}$ (Fig. 1 E, Supplementary Figure 2 A).

###### 3.4.2 Cellular volume

For computation of volume and membrane constraints we also needed to know what the actual cell volume is. We found no relation between growth rate and cell volume (Supplementary Figure 5). Therefore, we used the mean cell volume of

$\overline{V}_{cell} = 22.7\ \mu m^3$ in further calculations. Contrary to *P. kluyveri*, *S. cerevisiae* shows a linear relationship between cell size and growth rate (Tyson et al., 1979) (Table 2). For more information on these constraints we kindly refer to Elsemman et al. (Elsemman et al., 2022).

###### 3.4.3 Mitochondrial volume

Mitochondrial volume constraint sets a limit to how much mitochondrial protein volume fits inside a single cell. For *S. cerevisiae* we fitted the available mitochondrial protein volume ($V_{mitochondrialprotein}$) to 0.7 $\mu m^3$ such that the critical dilution rate for ethanol formation is at 0.28 $h^{-1}$ with this constraint active. For pcPichia we derived a mitochondrial volume constraint from $V_{mitochondrial,protein}$ of *S. cerevisiae*. With an absolute mitochondrial content of 0.7 $\mu m^3$ for *S. cerevisiae* and a cell volume of 28.8 $\mu m^3$ (Table 2), the mitochondrial protein volume occupies 2.43 % of the cellular volume ($\psi_{mitochondrialprotein}$). If *P. kluyveri* would have the same volume fraction, with mean cell volume of 22.7 $\mu m^3$ (Supplementary Figure 5) the mitochondrial volume constraint would be 0.552 $\mu m^3$. We did find mitochondrial mass fractions to be comparable when respiratory (Fig. 2 A).

However, based on available volume data, total mitochondrial volume of both species appear to differ: for *P. kluyveri* we measured an average mitochondrial volume fraction ($\psi_{mitochondria}$) of 10.6 % (Fig. 2 C), while $\psi_{mitochondria}$ of *S. cerevisiae* lies between 6.4 % in ethanol batch and 7.4 % in glucose batch conditions(Visser et al., 1995) (Table 2). To correct for this difference we calculated conversion factor alpha, using the highest mitochondrial content measurement of 7.4 % (Eq. 8).

$$\alpha = \frac{\psi^{S.cerevisiae}_{mitochondria}}{\psi^{P.klyuveri}_{mitochondria}} \tag{8}$$

Using factor $\alpha$ of 0.7, we can now calculate the relative mitochondrial protein volume in *P. kluyveri*, resulting in 3.48 % (Eq. 9).

$$V_{mitochondrialprotein} = \frac{V^{uncorrected}_{mitochondrialprotein}}{\alpha} \tag{9}$$

This results in an estimated absolute mitochondrial protein volume of $0.791 \mu m^3$.

Alex Bateman, Maria-Jesus Martin, Sandra Orchard, Michele Magrane, Rahat Agivetova, Shadab Ahmad, Emanuele Alpi, Emily H Bowler-Barnett, Ramona Britto, Borisas Bursteinas, Hema Bye-A-Jee, Ray Coetzee, Austra Cukura, Alan Da Silva, Paul Denny, Tunca Dogan, ThankGod Ebenezer, Jun Fan, Leyla Garcia Castro, Penelope Garmiri, George Georghiou, Leonardo Gonzales, Emma Hatton-Ellis, Abdulrahman Hussein, Alexandr Ignatchenko, Giuseppe Insana, Rizwan Ishtiaq, Petteri Jokinen, Vishal Joshi, Dushyanth Jyothi, Antonia Lock, Rodrigo Lopez, Aurelien Luciani, Jie Luo, Yvonne Lussi, Alistair MacDougall, Fabio Madeira, Mahdi Mahmoudy, Manuela Menchi, Alok Mishra, Katie Moulang, Andrew Nightingale, Carla Susana Oliveira, Sangya Pundir, Guoying Qi, Shriya Raj, Daniel Rice, Milagros Rodriguez Lopez, Rabie Saidi, Joseph Sampson, Tony Sawford, Elena Speretta, Edward Turner, Nidhi Tyagi, Preethi Vasudev, Vladimir Volynkin, Kate Warner, Xavier Watkins, Rossana Zaru, Hermann Zellner, Alan Bridge, Sylvain Poux, Nicole Redaschi, Lucila Aimo, Ghislaine Argoud-Puy, Andrea Auchincloss, Kristian Axelsen, Parit Bansal, Delphine Baratin, Marie-Claude Blatter, Jerven Bolleman, Emmanuel Boutet, Lionel Breuza, Cristina Casals-Casas, Edouard de Castro, Kamal Chikh Echioukh, Elisabeth Coudert, Beatrice Cuche, Mikael Doche, Dolnide Dornevil, Anne Estreicher, Maria Livia Famiglietti, Marc Feuermann, Elisabeth Gasteiger, Sebastien Gehant, Vivienne Gerritsen, Arnaud Gos, Nadine Gruaz-Gumowski, Ursula Hinz, Chantal Hulo, Nevila Hyka-Nouspikel, Florence Jungo, Guillaume Keller, Arnaud Kerhornou, Vicente Lara, Philippe Le Mercier, Damien Lieberherr, Thierry Lombardot, Xavier Martin, Patrick Masson, Anne Morgat, Teresa Batista Neto, Salvo Paesano, Ivo Pedruzzi, Sandrine Pilbout, Lucille Pourcel, Monica Pozzato, Manuela Pruess, Catherine Rivoire, Christian Sigrist, Karin Sonesson, Andre Stutz, Shyamala Sundaram, Michael Tognolli, Laure Verbregue, Cathy H Wu, Cecilia N Arighi, Leslie Arminski, Chuming Chen, Yongxing Chen, John S Garavelli, Hongzhan Huang, Kati Laiho, Peter McGarvey, Darren A Natale, Karen Ross, C R Vinayaka, Qinghua Wang, Yuqi Wang, Lai-Su Yeh, Jian Zhang, Patrick Ruch, and Douglas Teodoro. UniProt: the universal protein knowledgebase in 2021. *Nucleic Acids Research*, 49(D1):D480–D489, November 2020. doi: 10.1093/nar/gkaa1100. URL `https://doi.org/10.1093/nar/gkaa1100`.

Evert Bosdriesz, Meike T. Wortel, Jurgen R. Haanstra, Marijke J. Wagner, Pilar de la Torre Cortés, and Bas Teusink. Low affinity uniporter carrier proteins can increase net substrate uptake rate by reducing efflux. *Scientific Reports*, 8(1), April 2018. doi: 10.1038/s41598-018-23528-7. URL `https://doi.org/10.1038/s41598-018-23528-7`.

Hannah R. Bridges, Ljuban Grgic, Michael E. Harbour, and Judy Hirst. The respiratory complexes i from the mitochondria of two pichia species. *Biochemical Journal*, 422(1):151–159, July 2009. doi: 10.1042/bj20090492. URL `https://doi.org/10.1042/bj20090492`.

André B. Canelas, Cor Ras, Angela ten Pierick, Walter M. van Gulik, and Joseph J. Heijnen. An in vivo data-driven framework for classification and quantification of enzyme kinetics and determination of apparent thermodynamic data. *Metabolic Engineering*, 13(3):294–306, May 2011. doi: 10.1016/j.ymben.2011.02.005. URL `https://doi.org/10.1016/j.ymben.2011.02.005`.

Ibrahim E. Elsemman, Angelica Rodriguez Prado, Pranas Grigaitis, Manuel Garcia Albornoz, Victoria Harman, Stephen W.

Holman, Johan van Heerden, Frank J. Bruggeman, Mark M. M. Bisschops, Nikolaus Sonnenschein, Simon Hubbard, Rob Beynon, Pascale Daran-Lapujade, Jens Nielsen, and Bas Teusink. Whole-cell modeling in yeast predicts compartment-specific proteome constraints that drive metabolic strategies. *Nature Communications*, 13(1), February 2022. doi: 10.1038/s41467-022-28467-6. URL `https://doi.org/10.1038/s41467-022-28467-6`.

Kathrin Förster, Paola Turina, Friedel Drepper, Wolfgang Haehnel, Susanne Fischer, Peter Gräber, and Jan Petersen. Proton transport coupled atp synthesis by the purified yeast h+atp synthase in proteoliposomes. *Biochimica et Biophysica Acta (BBA) - Bioenergetics*, 1797(11):1828–1837, November 2010. doi: 10.1016/j.bbabio.2010.07.013. URL `https://doi.org/10.1016/j.bbabio.2010.07.013`.

Pranas Grigaitis, Samira L van den Bogaard, and Bas Teusink. Elevated energy costs of biomass production in mitochondrial respiration-deficient saccharomyces cerevisiae. *FEMS Yeast Research*, 23, 2023. doi: 10.1093/femsyr/foad008. URL `https://doi.org/10.1093/femsyr/foad008`.

Gary W. Grimes, Henry R. Mahler, and Philip S. Perlman. NUCLEAR GENE DOSAGE EFFECTS ON MITOCHONDRIAL MASS AND DNA. *The Journal of Cell Biology*, 61(3):565–574, June 1974. doi: 10.1083/jcb.61.3.565. URL `https://doi.org/10.1083/jcb.61.3.565`.

Ville R. I. Kaila and Mårten Wikström. Architecture of bacterial respiratory chains. *Nature Reviews Microbiology*, 19(5):319–330, January 2021. doi: 10.1038/s41579-020-00486-4. URL `https://doi.org/10.1038/s41579-020-00486-4`.

Eduard J. Kerkhoven, Kyle R. Pomraning, Scott E. Baker, and Jens Nielsen. Regulation of amino-acid metabolism controls flux to lipid accumulation in yarrowia lipolytica. *npj Systems Biology and Applications*, 2(1):16005, Mar 2016. ISSN 2056-7189. doi: 10.1038/npjsba.2016.5. URL `https://doi.org/10.1038/npjsba.2016.5`.

Petri-Jaan Lahtvee, Benjamín J. Sánchez, Agata Smialowska, Sergo Kasvandik, Ibrahim E. Elsemman, Francesco Gatto, and Jens Nielsen. Absolute quantification of protein and mrna abundances demonstrate variability in gene-specific translation efficiency in yeast. *Cell Systems*, 4(5):495–504.e5, May 2017. ISSN 2405-4712. doi: 10.1016/j.cels.2017.03.003. URL `http://dx.doi.org/10.1016/j.cels.2017.03.003`.

Hongzhong Lu, Feiran Li, Benjamín J. Sánchez, Zhengming Zhu, Gang Li, Iván Domenzain, Simonas Marcišauskas, Petre Mihail Anton, Dimitra Lappa, Christian Lieven, Moritz Emanuel Beber, Nikolaus Sonnenschein, Eduard J. Kerkhoven, and Jens Nielsen. A consensus s. cerevisiae metabolic model yeast8 and its ecosystem for comprehensively probing cellular metabolism. *Nature Communications*, 10(1):3586, 08 2019. ISSN 2041-1723. doi: 10.1038/s41467-019-11581-3. URL `https://doi.org/10.1038/s41467-019-11581-3`.

Simonas Marcišauskas, Boyang Ji, and Jens Nielsen. Reconstruction and analysis of a kluyveromyces marxianus genome-scale metabolic model. *BMC Bioinformatics*, 20(1):551, Nov 2019. ISSN 1471-2105. doi: 10.1186/s12859-019-3134-5. URL https://doi.org/10.1186/s12859-019-3134-5.

Avlant Nilsson and Jens Nielsen. Metabolic trade-offs in yeast are caused by f1f0-atp synthase. *Scientific Reports*, 6(1), March 2016. ISSN 2045-2322. doi: 10.1038/srep22264. URL http://dx.doi.org/10.1038/srep22264.

K. P. O'Brien. Inparanoid: a comprehensive database of eukaryotic orthologs. *Nucleic Acids Research*, 33(Database issue): D476–D480, December 2004. doi: 10.1093/nar/gki107. URL https://doi.org/10.1093/nar/gki107.

Brett Olivier. Systemsbioinformatics/cbmpy-metadraft: Metadraft is now available, 2018. URL https://zenodo.org/record/2398336.

Jan Petersen, Kathrin Förster, Paola Turina, and Peter Gräber. Comparison of the h+/atp ratios of the h+atp synthases from yeast and from chloroplast. *Proceedings of the National Academy of Sciences*, 109(28):11150–11155, June 2012. doi: 10.1073/pnas.1202799109. URL https://doi.org/10.1073/pnas.1202799109.

H. Schagger. Supercomplexes in the respiratory chains of yeast and mammalian mitochondria. *The EMBO Journal*, 19(8): 1777–1783, April 2000. doi: 10.1093/emboj/19.8.1777. URL https://doi.org/10.1093/emboj/19.8.1777.

Puneet Sharma, Benedikt S. Nilges, Jie Wu, and Sebastian A. Leidel. The translation inhibitor cycloheximide affects ribosome profiling data in a species-specific manner. *bioRxiv*, August 2019. doi: 10.1101/746255. URL https://doi.org/10.1101/746255.

Vincent Somerville, Pranas Grigaitis, Julius Battjes, Francesco Moro, and Bas Teusink. Use and limitations of genome-scale metabolic models in food microbiology. *Current Opinion in Food Science*, 43:225–231, February 2022. doi: 10.1016/j.cofs.2021.12.010. URL https://doi.org/10.1016/j.cofs.2021.12.010.

Patrick F. Suthers, Hoang V. Dinh, Zia Fatma, Yihui Shen, Siu Hung Joshua Chan, Joshua D. Rabinowitz, Huimin Zhao, and Costas D. Maranas. Genome-scale metabolic reconstruction of the non-model yeast issatchenkia orientalis sd108 and its application to organic acids production. *Metabolic Engineering Communications*, 11:e00148, 2020. ISSN 2214-0301. doi: https://doi.org/10.1016/j.mec.2020.e00148. URL https://www.sciencedirect.com/science/article/pii/S2214030120300481.

Màrius Tomàs-Gamisans, Pau Ferrer, and Joan Albiol. Fine-tuning the p. pastoris iMT1026 genome-scale metabolic model for improved prediction of growth on methanol or glycerol as sole carbon sources. *Microbial Biotechnology*, 11(1):224–237, November 2017. doi: 10.1111/1751-7915.12871. URL https://doi.org/10.1111/1751-7915.12871.

Tatsuhisa Tsuboi, Matheus P Viana, Fan Xu, Jingwen Yu, Raghav Chanchani, Ximena G Arceo, Evelina Tutucci, Joonhyuk Choi, Yang S Chen, Robert H Singer, Susanne M Rafelski, and Brian M Zid. Mitochondrial volume fraction and translation duration impact mitochondrial mRNA localization and protein synthesis. *eLife*, 9, August 2020. doi: 10.7554/elife.57814. URL `https://doi.org/10.7554/elife.57814`.

C B Tyson, P G Lord, and A E Wheals. Dependency of size of saccharomyces cerevisiae cells on growth rate. *Journal of Bacteriology*, 138(1):92–98, April 1979. ISSN 1098-5530. doi: 10.1128/jb.138.1.92-98.1979. URL `http://dx.doi.org/10.1128/jb.138.1.92-98.1979`.

H. Van Urk, E. Postma, W. A. Scheffers, and J. P. Van Dijken. Glucose transport in crabtree-positive and crabtree-negative yeasts. *Microbiology*, 135(9):2399–2406, September 1989. doi: 10.1099/00221287-135-9-2399. URL `https://doi.org/10.1099/00221287-135-9-2399`.

Pim van Hoek, Johannes P. van Dijken, and Jack T. Pronk. Regulation of fermentative capacity and levels of glycolytic enzymes in chemostat cultures of saccharomyces cerevisiae. *Enzyme and Microbial Technology*, 26(9-10):724–736, June 2000. doi: 10.1016/s0141-0229(00)00164-2. URL `https://doi.org/10.1016/s0141-0229(00)00164-2`.

Jean Velours and Geneviéve Arselin. The saccharomyces cerevisiae atp synthase. *Journal of Bioenergetics and Biomembranes*, 32(4):383–390, 2000. doi: 10.1023/a:1005580020547. URL `https://doi.org/10.1023/a:1005580020547`.

C. Verduyn, E. Postma, W. A. Scheffers, and J. P. van Dijken. Physiology of saccharomyces cerevisiae in anaerobic glucose-limited chemostat culturesx. *Journal of General Microbiology*, 136(3):395–403, March 1990. doi: 10.1099/00221287-136-3-395. URL `https://doi.org/10.1099/00221287-136-3-395`.

Wiebe Visser, Edwin A. van Spronsen, Nanne Nanninga, Jack T. Pronk, J. Gijs Kuenen, and Johannes P. van Dijken. Effects of growth conditions on mitochondrial morphology in saccharomyces cerevisiae. *Antonie van Leeuwenhoek*, 67(3):243–253, September 1995. doi: 10.1007/bf00873688. URL `https://doi.org/10.1007/bf00873688`.
